# Histology-based average template of the marmoset cortex with probabilistic localization of cytoarchitectural areas

**DOI:** 10.1101/2020.04.10.036632

**Authors:** Piotr Majka, Sylwia Bednarek, Jonathan M. Chan, Natalia Jermakow, Cirong Liu, Gabriela Saworska, Katrina H. Worthy, Afonso C. Silva, Daniel K. Wójcik, Marcello G.P. Rosa

**Affiliations:** Laboratory of Neuroinformatics, Nencki Institute of Experimental Biology of the Polish Academy of Sciences, 02-093 Warsaw, Poland; Australian Research Council, Centre of Excellence for Integrative Brain Function, Monash University Node, Clayton, VIC 3800, Australia; Biomedicine Discovery Institute and Department of Physiology, Monash University, Clayton, VIC 3800, Australia; Department of Neurobiology, University of Pittsburgh Brain Institute, Pittsburgh, PA, USA; Institute of Applied Psychology, Faculty of Management and Social Communication, Jagiellonian University, 30-348 Cracow, Poland

**Keywords:** Non-human primate, cortex, digital template, Nissl, cortical thickness, *Callithrix jacchus*

## Abstract

The rapid adoption of marmosets in neuroscience has created a demand for three dimensional (3D) atlases of the brain of this species to facilitate data integration in a common reference space. We report on a new open access template of the marmoset cortex (the Nencki–Monash, or NM template), representing a morphological average of 20 brains of young adult individuals, obtained by 3D reconstructions generated from Nissl-stained serial sections. The method used to generate the template takes into account morphological features of the individual brains, as well as the borders of clearly defined cytoarchitectural areas. This has resulted in a resource which allows direct estimates of the most likely coordinates of each cortical area, as well as quantification of the margins of error involved in assigning voxels to areas, and preserves quantitative information about the laminar structure of the cortex. We provide spatial transformations between the NM and other available marmoset brain templates, thus enabling integration with magnetic resonance imaging (MRI) and tracer-based connectivity data. The NM template combines some of the main advantages of histology-based atlases (e.g. information about the cytoarchitectural structure) with features more commonly associated with MRI-based templates (isotropic nature of the dataset, and probabilistic analyses). The underlying workflow may be found useful in the future development of brain atlases that incorporate information about the variability of areas in species for which it may be impractical to ensure homogeneity of the sample in terms of age, sex and genetic background.

**Graphical abstract:** 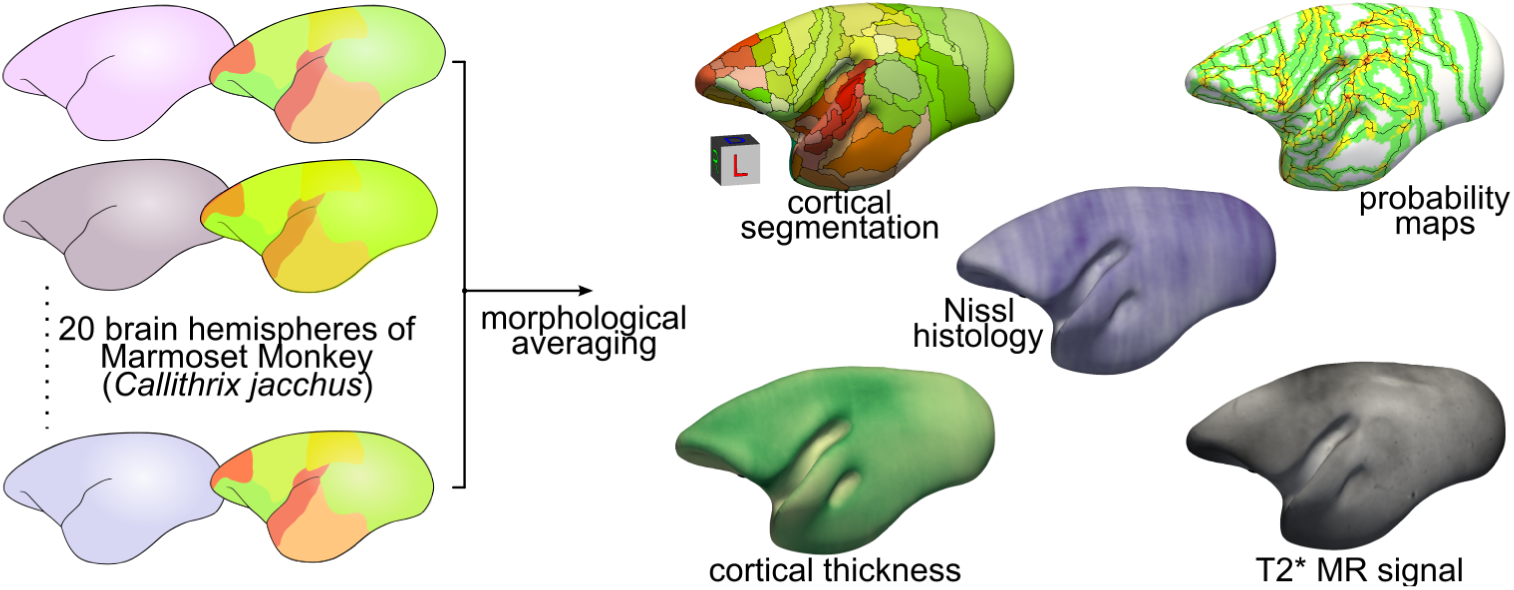

**Highlights:**

- A 3D template of the marmoset cortex representing the average of 20 individuals.
- The template is based on Nissl stain and preserves information about cortical layers.
- Probabilistic mapping of areas, cortical thickness, and layer intensity profiles.
- Includes spatial transformations to other marmoset brain atlases.

**Abbreviations:** For a list of areas and their abbreviations see Table S2.

## 1. Introduction

Brain templates are a key resource for current efforts towards brain mapping, which allow neuroinformatics approaches to data analysis and integration across modalities (Evans et al., 2012). The importance of templates that accurately represent the brain in 3 dimensions has been demonstrated by their widespread adoption in large–scale projects for studies of the human (Shattuck et al., 2008; Glasser et al., 2016), macaque (Feng et al., 2017; Reveley et al., 2017; Seidlitz et al., 2018) and rodent (Schweinhardt et al., 2003; Kuan et al., 2015) brains, among other species. This has resulted in marked progress in our understanding of the relationship between brain structure and function, as well as of the changes associated with pathological conditions, development, and brain evolution (Van Essen and Glasser, 2018).

The marmoset (*Callithrix jacchus*) is a rapidly emerging non-human primate model with complementary advantages to those offered by the traditionally used macaque monkey. In particular, marmosets offer benefits in terms of a faster developmental cycle, which facilitates both developmental studies and the generation of transgenic lines (Tomioka et al., 2017; Okano and Kishi, 2018; Sawiak et al., 2018). The relatively simpler brain morphology also enables a wider variety of physiological studies involving multielectrode arrays (Zavitz et al., 2016; Ghahremani et al., 2019), optogenetics (MacDougall et al., 2016; Nurminen et al., 2018), and 2-photon imaging (Ebina et al., 2018; Zeng et al., 2019). At the same time, the marmoset cortex contains sophisticated neural systems associated with vision and hearing which characterize primate brains, as well as many prefrontal, premotor, posterior parietal and temporal association areas which are not clearly differentiated in rodents (Solomon and Rosa, 2014; Bakola et al., 2015; Miller et al., 2016; Majka et al., 2020).

Presently, there are several digital templates of the marmoset brain, which have been successfully used in a variety of physiological and anatomical studies (e.g. Toarmino et al., 2017; Atapour et al., 2019; Ghahremani et al., 2019; Hori et al., 2020; Liu et al., 2019; Majka et al., 2019, 2020). However, most atlases are based on single individuals, with the brains processed using histological techniques (Majka et al., 2016; Woodward et al., 2018) or imaged with high field magnetic resonance imaging (MRI; Liu et al., 2018). All of these templates have adopted the scheme of parcellation first proposed by Paxinos et al. (2012). More recently, a population template based on 3T MR images of 13 individuals was introduced (Risser et al., 2019), but it was also based on the registration of the same specimen studied by Paxinos et al. (2012) to multiple brains. As discussed recently (Majka et al., 2020), there is a significant variation in the cortex between individuals, which results in potential errors in the assignment of voxels to areas in studies involving the integration of data from multiple individuals.

Here we introduce a template of the cerebral cortex resulting from the morphological averaging of 20 three-dimensional (3D) reconstructions of young adult marmoset brains (the Nencki–Monash, or NM template). Unlike previously available atlases, the present resource is based on the analysis of cytoarchitectural borders in multiple individuals, visualized in Nissl-stained coronal sections. Among other innovations, this approach allows probabilistic allocation of voxels to cortical areas, and quantification of the anatomical variability across the brain. In addition, the template generation procedure (Avants et al., 2011) emphasizes the common cytoarchitectonic features between individual brains while mitigating the impact of histology artifacts. The NM atlas is distributed under an open license and includes spatial transformations required to align the template to other currently available atlases of the marmoset brain, thus enabling future data integration across laboratories. We, therefore, consider the Nencki–Monash template a significant step towards data federation for large-scale projects based on marmosets.

## 2. Material and methods

### 2.1 Data selection and quality assessment

The Nencki-Monash template is based on 20 three-dimensional reconstructions of single hemispheres of marmoset brains, selected among the cases available in the Marmoset Brain Connectivity Atlas portal (http://marmosetbrain.org). These brains were originally a part of a series of studies involving injections of retrograde tracers in the cortex; for a comprehensive description of the histological procedures, the reader is referred to the original papers in this series of studies (e.g. Burman et al., 2014a,b; Majka et al., 2020, http://www.marmosetbrain.org/reference). In all the cases, 40 µm thick frozen sections were obtained from paraformaldehyde-fixed brains. One in 5 sections was set aside for Nissl staining (with cresyl violet), and two adjacent series were stained for myelin and cytochrome oxidase; the two remaining series were used for analysis of connections using fluorescent tracers, and immunocytochemical procedures (which varied between cases, see e.g. Atapour et al., 2019). Although all histological series contributed to the parcellation of the cortex into areas, the template was generated based on the Nissl-stained series (see also Majka et al., 2020).

The pool of cases was initially narrowed down to those sectioned in the coronal plane. Cases with faint, highly uneven, or otherwise ill stained sections were subsequently excluded, as were those with lesions or significant surgery-related distortions in the cortex, and those in which the sections did not cover the entire rostrocaudal extent of the brain. A margin of approximately 600 μm (equivalent to three missing sections, in a 1-in-5 series) was allowed at the frontal and occipital poles, as these sections are frequently too small and were not consistently collected in a typical sectioning process. The remaining cases were inspected for distortions and damages to the sections, and those with significant tears, stretches, frequently missing parts of the cortex were discarded. Minor imperfections inherent to histological procedures were allowed, provided they appeared sporadically in a given reconstruction. Finally, the individual 3D reconstructions themselves had to be of a sufficiently high quality, according to criteria that included no sharp transitions between sections or jagged edges, and evidence of smooth and clearly distinguishable anatomical details throughout. Ultimately, twenty most suitable young adult cases (1.4–4.6 years, median age: 2.8 years; 11 males) were selected for the template generation. The weight of the animals ranged from 304 g (female, 1.6 years) to 530 g (female, 4.7 years). Detailed metadata for each animal are available in Table 1.

**Table 1:** List of the brain hemispheres selected for the construction of the Nencki-Monash template.

| Case, hemisphere | Sex | Weight (g) | Age upon perfusion | Number of outlined areas |
| --- | --- | --- | --- | --- |
| CJ123, R | F | 530 | 4 years, 8 months | 19 |
| CJ114, R | M | 409 | 3 years, 4 months | 19 |
| CJ115, R | F | 387 | 2 years, 11 months | 18 |
| CJ116, R | F | 339 | 3 years, 2 months | 18 |
| CJ153, L | F | 336 | 1 year, 7 months | 18 |
| CJ174, R | F | 342 | 2 years, 8 months | 18 |
| CJ111, L | M | 334 | 3 years, 1 month | 18 |
| CJ108, R | M | 349 | 3 years, 2 months | 17 |
| CJ102, L | F | 350 | 4 year, 12 days | 17 |
| CJ56, L | M | 350 | 1 year, 8 months | 17 |
| CJ180, R | M | 374 | 2 years, 9 months | 17 |
| CJ112, L | M | 377 | 3 years, 2 months | 16 |
| CJ157, L | F | 364 | 1 year, 7 months | 16 |
| CJ164, L | F | 304 | 1 year, 7 months | 15 |
| CJ110, L | M | 357 | 3 years, 1 month | 15 |
| CJ81, L | F | 419 | 2 years, 5 months | 15 |
| CJ71, R | M | 412 | 1 year, 6 months | 15 |
| CJ122, R | M | 318 | 2 years, 11 months | 15 |
| CJ78, L | M | 394 | 2 years, 1 month | 13 |
| CJ181, R | M | 368 | 2 years, 3 months | 11 |

### 2.2 Input data preprocessing

The process of 3D reconstruction of individual hemispheres was described in detail in Majka et al. (2016, 2020); here, we use the direct output of these procedures as a substrate for the Nencki-Monash template generation. The input 3D reconstructions comprised between 144 (case CJ116) and 160 (case CJ181) sections (153 on average) spaced equally every 200 μm (see Majka et al., 2016), which resulted in a 3D NIfTI image (https://nifti.nimh.nih.gov/nifti-1) of a 40 × 200 × 40 μm (mediolateral, rostrocaudal and dorsoventral directions, respectively) resolution. Slight, but inevitable variation in staining intensity between the sections was mitigated with the use of the weighted histogram matching approach (Li et al., 2009). In each stack, sections standing out in terms of intensity (i.e., noticeably darker or lighter than adjacent sections) were identified. Subsequently, the intensity distribution of each channel of the RGB section image was, independently, matched with the one of an average image of adjacent sections resulting in an adjusted image that replaced the initial one.

### 2.3 Mapping the parcellation from the Paxinos et al. (2012) template onto individual cases

The cerebral cortex was outlined manually in each of the reconstructions using ITK-SNAP software (http://www.itksnap.org, Yushkevich et al., 2006), in a manner that covered all isocortical areas, as well as the pericallosal, piriform, insular, parahippocampal and entorhinal complexes, but excluding the hippocampal formation. In the next step, the cortex was segmented into 116 areas, according to the Paxinos et al. (2012) marmoset brain atlas parcellation (Fig. 1). This was accomplished by registering the individual cases to the three-dimensional version of the atlas available at http://www.marmosetbrain.org/reference.

**Figure 1:**
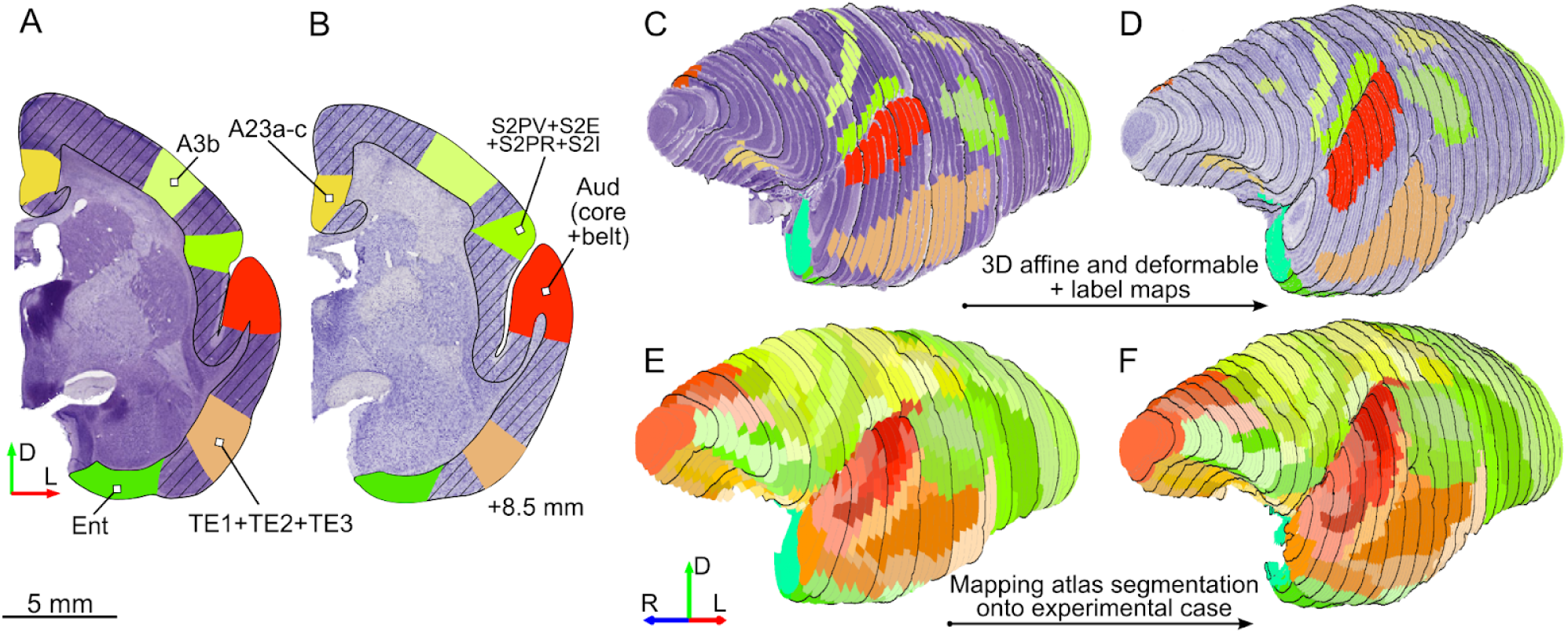
Registration-based segmentation of an individual case into cortical areas. Label-guided segmentation of individual hemispheres into cortical areas illustrated using an example case (CJ116). Different colors represent cytoarchitectural areas delineated manually to guide the coregistration with Paxinos et al. (2012) atlas. For detailed information on areas outlined in each hemisphere see Table S1. **A, B**: Example coronal sections (8.5 mm rostral to the interaural line) from the Paxinos et al. (2012) atlas (A) and the experimental (B) brain with a set of corresponding label maps indicated. The cerebral cortex was outlined (diagonal hatching) as an additional constraint. **C-F**: Rostrolateral views of reconstructions of the marmoset brain. Black outlines on every fifth section are provided to better illustrate the general shape of the image stack and are unrelated to the mapping process. **C**: reconstruction of the brain illustrated in the Paxinos et al. (2012) atlas. **D**: reconstruction of the experimental case. **E**: Segmentation of the Paxinos et al. (2012) atlas into cortical areas. **F**: Segmentation from panel E mapped onto the experimental case.

The registration procedure differed from typical approaches as, apart from the usual intensity-based registration, our solution also involved sets of label maps (corresponding to cytoarchitectural areas) delineated manually to increase the mapping accuracy (Majka et al., 2020, Fig. 1A, B). Depending on the case, between 11 and 19 areas were manually delineated by an expert anatomist (MGPR). Among these, a collection of eight label maps was consistently drawn in each hemisphere: area 3b (A3b), combined auditory core and belt areas, entorhinal cortex (Ent), piriform cortex (Pir), prostriate area (ProS), temporal area TH, primary visual cortex (V1), and middle temporal visual area (MT). These and other outlined areas were evenly distributed across the cortex (Fig. S1, see Table S1 for detailed information about the areas delineated in each hemisphere), with a slightly higher concentration in the regions which, according to our experience, pose more challenges in registration (Majka et al., 2020). The process of delineation always took into consideration the information provided by the interleaved myelin, and cytochrome oxidase-stained sections, as well as the Nissl-stained sections.

Each hemisphere was then independently mapped to the Paxinos et al. (2012) template (Fig. 1C, D), which represents the right hemisphere; where necessary, left hemisphere reconstructions were flipped horizontally. Before the registration, the 3D images representing both the template and the brains of individual animals were resampled to an isotropic resolution of 75 µm and smoothed with a median filter. The subsequent affine alignment was driven by Mattes mutual information (MI, Mattes et al., 2001) image similarity metric. The non-linear registration utilized the Symmetric Normalization algorithm SyN (Avants et al., 2011) and relied on two, equally weighted, metrics. For the intensity images (i.e. Nissl stain), the normalized correlation coefficient (CC, Avants et al., 2008) was used. The window size was set to 5 voxels with the gradient step of 0.25; the velocity field was regularized with a Gaussian kernel (standard deviation of one voxel) while the smoothing of the displacement field was switched off. Simultaneously, the overlap between corresponding label maps (Fig. 1C, D) was enforced by the use of Point-Set Estimation (PSE, Avants et al., 2011) metric with an exhaustive (100%) sampling of the labelled voxels. This process established a mapping between an experimental case and the Paxinos et al. (2012) template taking into account constraints imposed by expert-outlined cortical areas. The computed transformation was then used to map the Paxinos et al. (2012) parcellation (Fig. 1E) of the cortex onto the experimental 3D reconstruction (Fig. 1F).

### 2.4 Average template construction

The overview of the process of generating the average histological volume of the Nencki-Monash template and auxiliary datasets is presented in Fig. 2. The histological volumes of individual marmoset brains (Fig. 1D) were first converted to grayscale by extracting the red image channel and then resampled to an isotropic resolution of 100 µm. The average morphology was computed using the Symmetric Groupwise Normalization formalism (SyGN, Tustison et al., 2014) implemented in the ANTS software suite (Avants et al., 2011, `buildtemplateparallel.sh` script). The initial target image was created by the affine alignment of the histological volumes to the Paxinos et al. (2012) atlas, using the transformations obtained as described above (Fig. 1C, D and Fig. 2A), followed by voxel-wise averaging of the transformed images. Each of the 20 iterations of the morphological averaging procedure (Fig. 2B) consisted of a nonlinear registration (SyN transformations, CC similarity metric, three-level image pyramid, per-iteration affine alignment disabled) of histological volumes to an intermediate template. Upon computing the spatial transformations, the initial 3D images were warped and averaged voxel-wise to obtain an updated, more precise target image. Eventually, the initial RGB histological volumes were transformed using the displacement fields computed in the final iteration and rendered on a 50 µm image grid. The resulting image (Fig. 2C) is not biased towards any of the individual histological volumes (Avants and Gee, 2004), and, by taking advantage of the subpixel accuracy of the morphological averaging (Janke et al., 2015), has a higher spatial resolution than the input 3D images. The same set of transformations was applied to the segmentations of the input brain hemispheres (Fig. 2D), and for each of the 116 areas, a map (Fig. 2E) was computed showing the probability of encountering a given cortical area in a selected voxel. Both the template image and the probability maps underwent streamline-based analysis (Fig. 2F) in order to calculate the segmentation into cortical areas (Fig. 2G) and auxiliary datasets (Fig. 2H-J).

**Figure 2:**
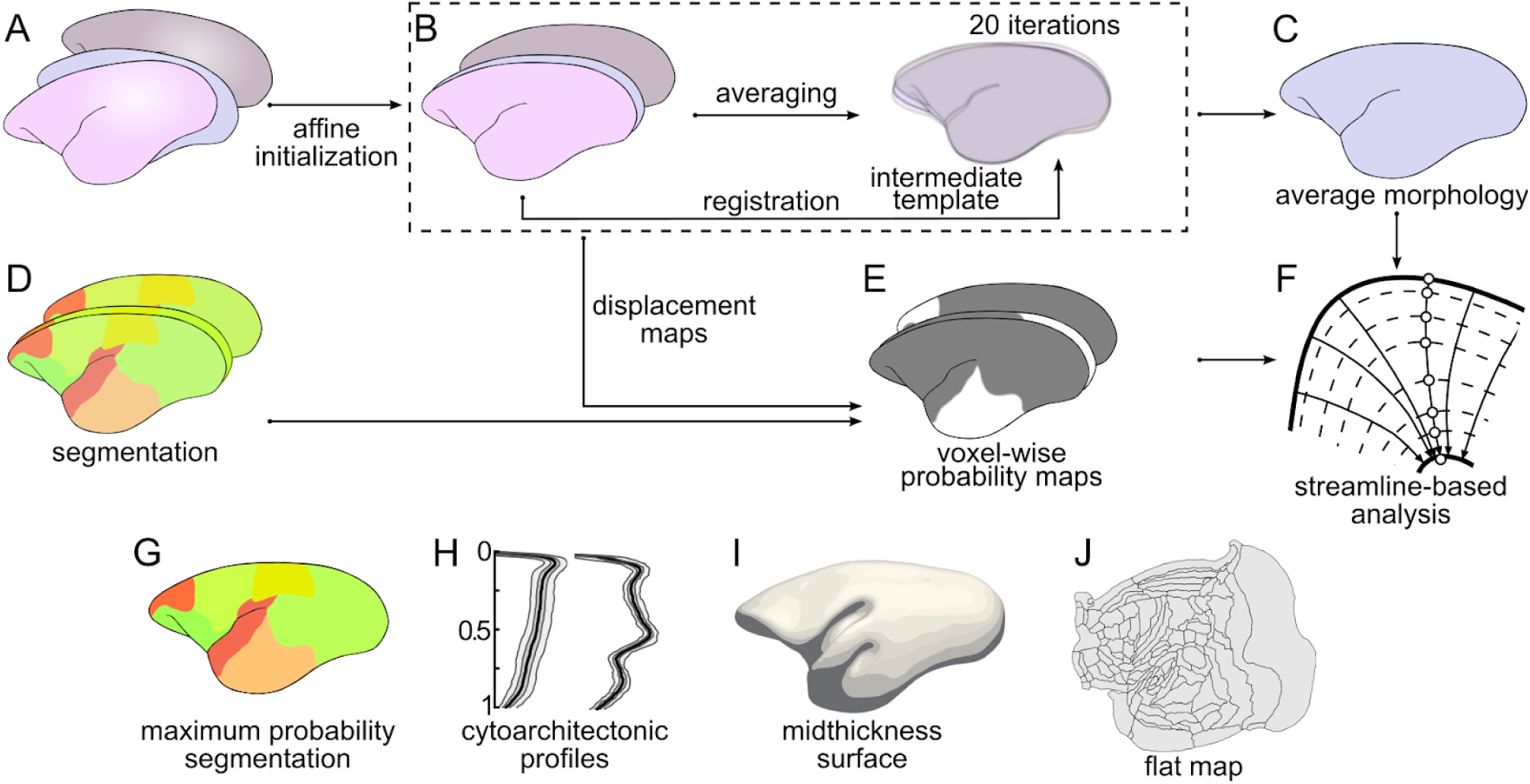
Nencki–Monash template generation pipeline. **A**: Set of 20 histological volumes constituting the input for the template generation (see Fig. 1B, D). Histological volumes affinely registered to Paxinos et al. (2012) marmoset brain atlas (**B**) and averaged voxel-wise constitute the starting point for the template generation. Next, 20 iterations of the Symmetric Groupwise Normalization (SyGN, Tustison et al., 2014) procedure are carried out (see text for details) resulting in the average histological volume (**C**). The segmentations of individual histological volumes (**D**) obtained using the procedure illustrated in Fig. 1F are transformed and probability maps (**E**) for each cortical area are calculated. These results allow then for the analyses of various quantities perpendicular to the cortical layers (**F**, see Fig. 3, and section *2*.*5 Streamline-based analyses*), which resulted in the maximum probability segmentation of the template (**G**) and the cytoarchitectonic profiles of individual cortical areas (**H**). The profile-based analysis also provides convenient means to visualize the data related to the template by displaying it against the mid-thickness surface (**I**) or by projecting onto a flat map (**J**).

### 2.5 Streamline-based analyses

To enable profile-based analyses (i.e. analyses relying on various quantities measured perpendicular to the cortical layers, Fig. 3) we employed a 2-surface Laplacian-based approach (Jones et al., 2000, see Fig. 3A). The calculations were performed using an in-house implementation of the method written in Python, and the Visualization Toolkit framework (Schroeder, 2006; http://www.vtk.org/). The objective of this procedure was to generate streamlines that accurately estimated the local orientation of cortical columns (see also Wagstyl et al., 2018, Atapour et al., 2019).

**Figure 3:**
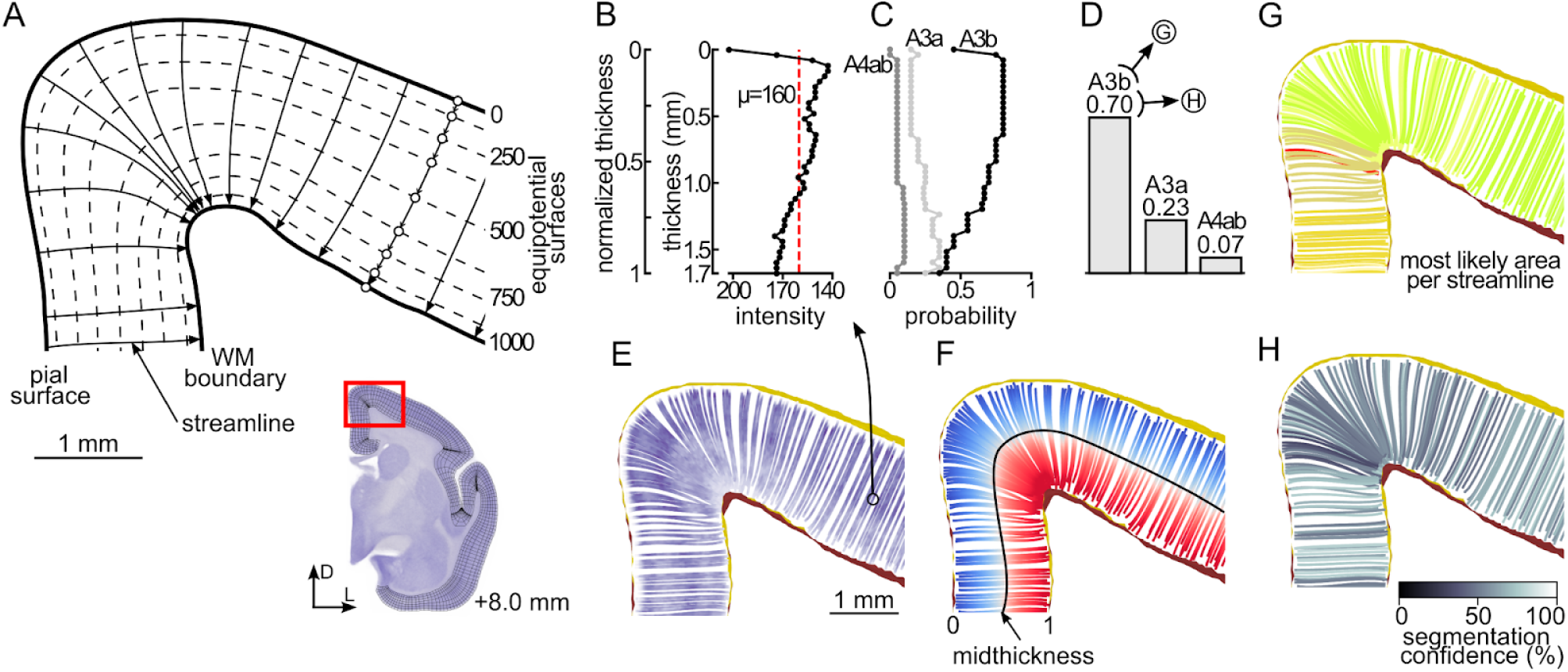
Overview of the streamline-based analyses. **A**: Schematic illustration of the 2-surface Laplacian-based method (Jones et al., 2000) using an example coronal cross-section located approximately 8.0 mm rostral from the interaural line. The drawing depicts the dorsomedial part of the section as indicated on the inset. The perimeter of the cortex is divided into the outer (the pial) and inner (the border between the gray and the white matter) surfaces and assigned with potential values (0 and +1000, respectively). The streamlines (see text for a detailed description of the process of computing the streamlines) start on the pial surface and traverse towards the white matter boundary intersecting the equipotential surfaces (dashed lines) at the right angle. **B**: Example laminar profile obtained by sampling the template at each point of the trajectory (white circles in panel A; for clarity only 9 out of 43 points are presented). The image intensity (values of the red channel of the template image) plotted against absolute and normalized thickness values. The average intensity and the thickness are then assigned to the streamline. **C**: Probabilities of encountering individual areas along the examined profile based on the voxel-wise probability maps (Fig. 2E). **D**: The probabilities are summed, normalized to one, and the streamline is labeled as belonging to the most likely area (G). Further, the highest total probability is expressed as percentages and referred as the segmentation confidence (H). **E**-**H**: Examples of various quantities examined in the profile-based analyses, for clarity, only approximately 5% of all profiles are shown: **E**: Image intensity, **F**: Normalized thickness; the points halfway across the profile (black line) are combined into the mid-thickness surface (see Fig. 2I). **G**: Most probable area per streamline; different colors denote various cortical areas according to the atlas (Fig. 1E). **H**: Segmentation confidence. For a list of areas and their abbreviations see Table S2.

The perimeter of the cortex was divided into the pial surface and the white matter (WM) boundary, and assigned with potential values (0 for the pial surface, +1000 for the WM boundary). Afterwards, a Laplacian equation was solved on a grid of one point per voxel, yielding the distribution of potential within the cortical mask. A gradient of the potential was computed producing a dense (defined for each voxel) vector field. Subsequently, 829 514 points constituting the pial surface were selected. Starting from each point, a trajectory (streamline) was computed by advancing a distance of 20 μm per iteration along a direction defined by the gradient vector until reaching the WM boundary. At each step, the following values were sampled: distance from the beginning of a streamline (an estimate of the depth below the pial surface, Fig. 3B), image intensity (Fig. 3B, E), probabilities of encountering each of the 116 areas (Fig. 3C) based on the previously computed maps (Fig. 2E); Upon reaching the WM border, the relative depth below the pial surface (Fig. 3F), the average intensity along the trajectory (Fig. 3B), and the most likely area based on the probabilities summed across all points of a streamline (Fig. 3D, G, and H) were determined. For additional analyses, the above mentioned values, recorded at individual points of the trajectories, were then projected onto the NM template image grid.

Next, a surface defined by points of normalized cortical thickness equal to 0.5 was extracted (the mid-thickness surface, Figs. 3F and 2I), and converted to a flat map (Fig. 2J) using an open-source, software package CARET (Computerized Anatomical Reconstruction and Editing Toolkit, Van Essen et al., 2001; http://brainvis.wustl.edu/wiki/index.php/Caret:About) and the approach described previously (Reser et al., 2013; Burman et al., 2015).

For the analyses relying on the cortical profiles, the streamlines with a high confidence level (90% or more, Fig. 3H) belonging to a chosen area were selected. When necessary, this confidence level threshold was gradually lowered by 5 percentage points in each step, until a minimum number of 200 streamlines was obtained. For the majority of the areas (88 out of 116), the initial threshold was sufficient, while the lowest confidence level necessary for gathering the required number of trajectories was 50% (areas APir and VIP). Once the streamlines for a given area were determined, the distribution of the length of the trajectories, as well as their average length, were calculated, with the latter interpreted as the mean cortical thickness of an area.

The same set of streamlines was used to calculate the staining intensity profiles (see Schleicher et al., 2000; Wagstyl et al., 2018). The Nissl-modality 3D image was sampled at 100 equidistant points along each streamline resulting in a set of profiles represented in the domain of the normalized thickness (0–pial, 1–WM boundary) to enable comparisons of the streamlines regardless of their absolute length. Based on that, profiles representing the lower and upper quartiles, median, as well as the 5th and 95th centiles of the intensities along streamlines selected for each area were defined (Fig. 2H).

### 2.6 Registration to other brain templates

To enable interoperability with other marmoset brain atlases, we registered the Nencki–Monash template to three other publicly available marmoset brain templates: the one based on the Paxinos et al. (2012) atlas of the marmoset brain (Majka et al., 2016), the template developed by the Brain/MINDS consortium (Woodward et al., 2018) and the Marmoset Brain Mapping (MBM) 2.0 marmoset brain template (Liu et al., 2020). All these templates were released with cortical parcellations that follow the Paxinos et al. (2012) convention, which makes it possible to augment the conventional intensity-based registration with the approach relying on label maps, analogously to the procedure demonstrated in Fig. 1.

Conceptually, the registration of the NM template to all three considered atlases was performed in a similar way but with some differences in the registration parameters (see Table S3). Initially, the target atlas image was masked so that only those parts of the brain tissue which have their counterparts in the NM template remained. This includes, for instance, removing the cerebellum and brainstem structures posterior to the caudal end of the thalamus. This step established the correspondence between the target atlas and the NM template, mitigating the chance of gross misregistration. The same mask was then applied to the label maps (segmentation into cortical areas). Initially, the images were aligned with the affine transformation, followed by a deformable registration step. The latter relied on simultaneous intensity- and label map-based registration which forced the algorithm to maximize the overlap of all of the 116 outlined cortical areas resulting in a precise mapping between the target atlas and the NM template. To evaluate the registration accuracy, the agreement between corresponding labels was assessed with the Dice and Jaccard coefficients as well as the Hausdorf distance using the `LabelOverlapMeasures` and the `HausdorffDistance` image filters implemented in the SimpleITK toolkit (http://www.simpleitk.org/).

## 3. Results

### 3.1 An average template of the marmoset brain

The Nencki-Monash template of the young adult marmoset brain, created from three-dimensional reconstructions of 20 individuals, is available for download, as a 50 µm isotropic NIfTI image, from http://www.marmosetbrain.org/nencki_monash_template. The template is embedded in a stereotaxic space compatible with the one used in the Paxinos et al. (2012) atlas, which relies on cranial landmarks: dorsoventral (DV) coordinates are relative to a horizontal zero plane that passes through the lower margin of the orbits and the center of the external auditory meatuses, anteroposterior (AP) coordinates are relative to a plane perpendicular to the horizontal zero and through the centers of the external auditory meatuses, and mediolateral (ML) coordinates are relative to the midsagittal plane. The template brain extends 31 mm along the AP axis, 11.8 mm along the ML axis, and 17.8 mm along the DV axis.

### 3.2 The template preserves information about the laminar structure of the cortex

The data layers associated with the template are summarized in Figures 4 and S2 (see Figures S3 and S4 for corresponding views in other cardinal planes). The process of generating the template preserved a significant amount of anatomical detail, sufficient to detect changes related to the expected laminar characteristics of the different cytoarchitectural areas, when visualized at low power. For example, the cortex assigned to dorsocaudal area 6 (A6DC) appears as nearly uniform in terms of gray level density across layers (Fig. 4A, A+12.3), in marked contrast with the characteristics of the primary somatosensory area (A3b), where a clear cell-dense band appears at the level of cortical layer 4, and there is a differentiation between a lightly stained layer 5 and a darker layer 6 (Fig. 4A, A+10.4). Further, isocortical area TE3 of the inferior temporal cortex shows obvious gray level differentiation associated with the 6 cortical layers, whereas the entorhinal cortex (Ent; comprising Brodmann’s areas 28 and 34) shows a clearly defined, lightly stained *lamina dissecans* (Ramón y Cajal, 1909) separating superficial and deep, darkly stained strata (corresponding to layers 2/3 and 5/6, respectively; Fig. 4A, A+7.4). These characteristics correspond to well-described cytoarchitectural features of these areas in the marmoset (Burman et al., 2008, 2011, 2014b; Góis Morais et al., 2019). Although subcortical structures were not analysed, the quality of the template also allows visualization of details such as the layers of the lateral geniculate nucleus (LGN) and other thalamic nuclei (Fig. 4A, A+4.4).

**Figure 4:**
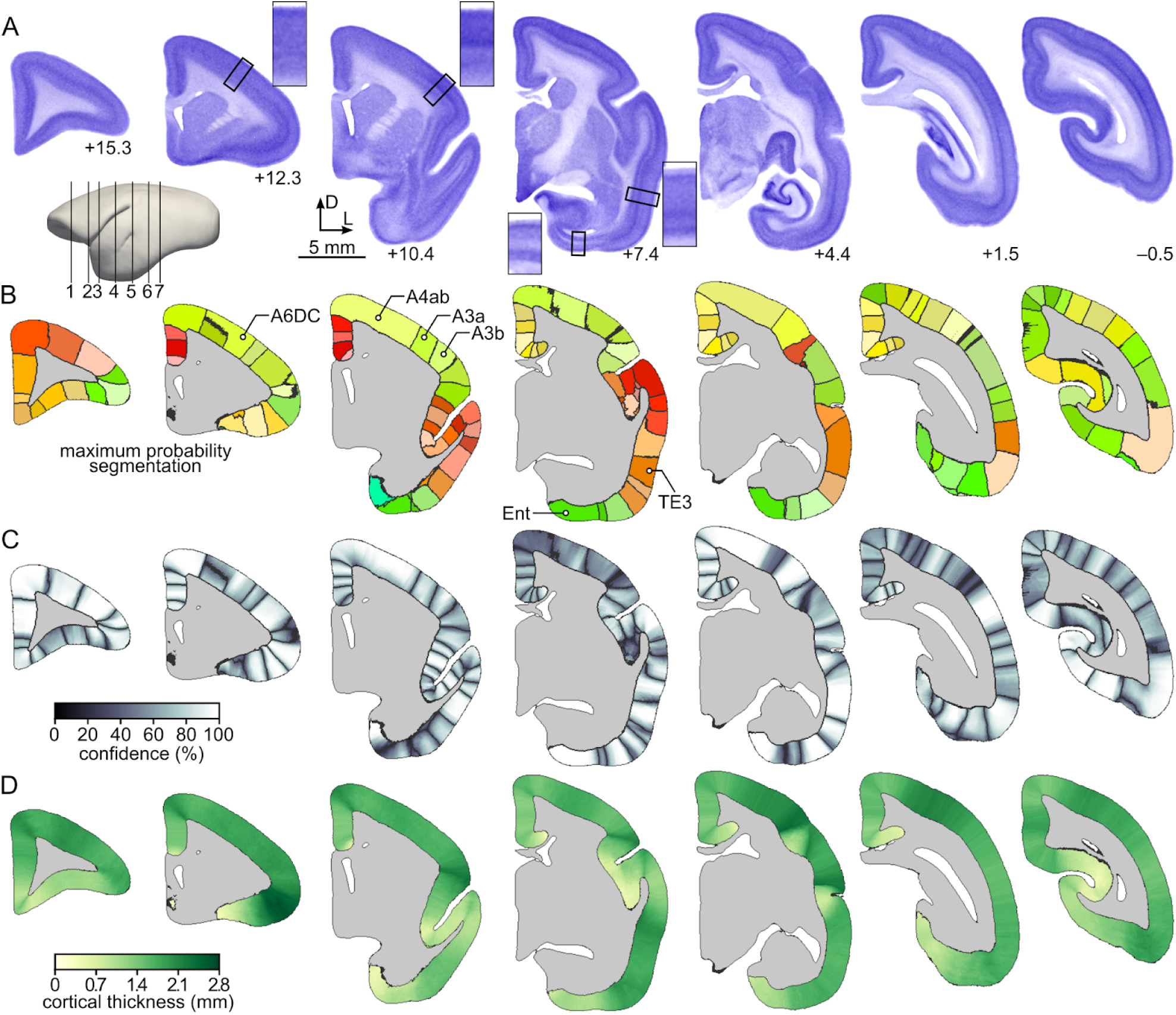
The main data layers of the Nencki–Monash template. Coronal cross-sections at various anterior-posterior levels illustrating the main layers of data constituting the template: **A**: The template represented as a Nissl-stained volume. A schematic lateral view of the template (left) indicates the levels of coronal sections corresponding to anteroposterior levels ranging between –0.5 and +15.3. The magnified regions next to some of the sections illustrate the laminar characteristics of selected cytoarchitectural areas labelled in panel B. **B**: Maximum probability segmentation of the cerebral cortex into 116 areas created by selecting, for each cortical profile, its most probable label (Fig. 3C, D). The black contours represent the borders of individual areas and are drawn for the clarity of the visualization. **C**: Segmentation confidence level map. **D**: Cortical thickness map. Additional two data layers of the template are presented in Fig. S2, while cross-sections in horizontal and parasagittal planes are shown in Figs. S3 and S4. All datasets are available for download from http://www.marmosetbrain.org/nencki_monash_template.

### 3.3 Boundaries of cortical areas and variability

Figure 5 summarizes the segmentation of the cortex according to the criteria and nomenclature established by Paxinos et al. (2012). The segmentation was obtained by selecting the most frequently encountered cortical area along a given streamline (Fig. 3C), thus incorporating the assumption that each cortical column belongs to a single area. This resulted in an anatomically plausible segmentation, with borders extending radially and crossing the laminar layers at the right angle (Fig. 5A).

**Figure 5:**
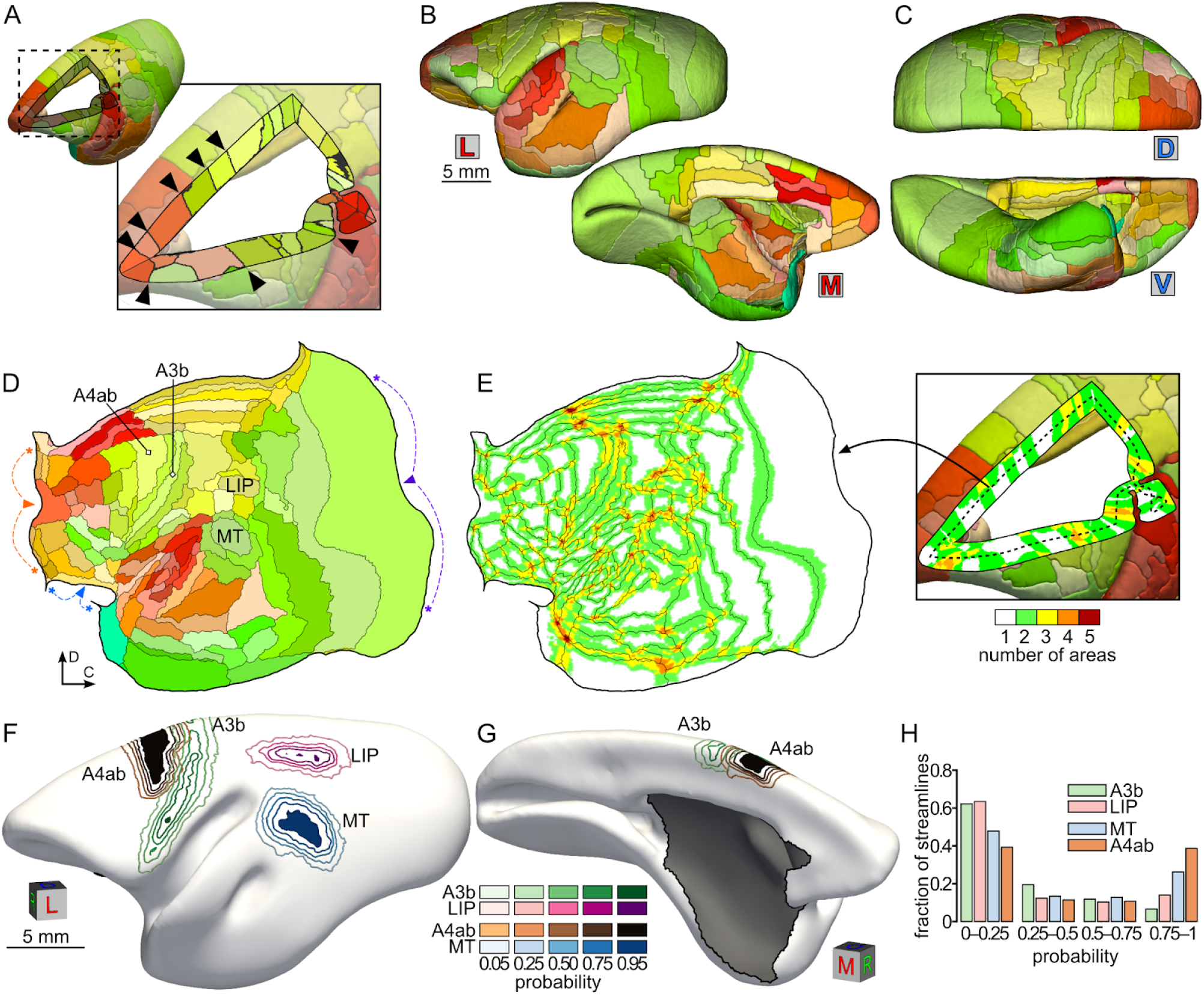
Segmentation of the Nencki–Monash template into cortical areas and analysis of variability of the areas’ location. **A**: Anatomically plausible properties of the segmentation relying on the most likely area along each cortical profile (black triangles). Cutout view through the frontal cortex, as indicated on the thumbnail 3D visualization above the panel. **B, C**: Medial (M), lateral (L), dorsal (D), and ventral (V) views of a 3D visualization of the template segmentation into 116 cortical areas, following the scheme proposed by Paxinos et. al. (2012). Colors denote different areas (see Table S2 for a list). **D**: Flat map representation of the segmentation shown on panels B-C. Arrowheads and asterisks indicate three points along the perimeter of the map where discontinuities were made to minimise distortions (fundus of the calcarine sulcus in purple, boundary between orbitofrontal cortex and medial frontal cortex in orange, piriform cortex in blue). **E**: Spatial variability of the location of cortical areas expressed as the numbers of areas with a probability ≥ 0.05 along each streamline. **F**-**G**: Analysis of the spatial variability of four example areas (A4ab, A3b, LIP, MT). The contours illustrated on rostrolateral (F) and rostromedial (G) views of the mid-thickness surface denote isoprobability lines (see color scales), and the dark-filled regions indicate parts of the cortex in which given area appears consistently (p≥ 0.95) throughout the sample. **H**: Quantitative analysis of the variability of the 4 example areas. The graph represents the fraction of streamlines with a non-zero probability of encountering a given area. Areas A3b and LIP express high spatial variability (higher fractions of low-probability streamlines) in comparison with areas A4ab and MT.

The process of segmentation of the individual hemispheres used to compute the NM template (Fig. 1) was guided by label maps corresponding to the actual extent of areas in each individual, based on the manual delineation of cyto-, myelo- and chemoarchitectural boundaries that could be unambiguously visualized. The incorporation of this procedure, which is unique to the present workflow, noticeably increased the precision of the delineation of cortical areas, as can be demonstrated by comparing how well the areas outlined manually match those of the Paxinos et al. (2012) atlas with and without using the label maps to drive the registration. (Fig. S5). As a result, the gross cytoarchitectural transitions that can be observed in the template (Fig. 4A) correlate well with the parcellation of the cortex (Fig. 4B, S3B and S4B), which takes into account variability across all 20 cases (Figs. 4C, 5B-D). For example, the average border between somatosensory areas 3b and 3a in Fig. 4A (+10.4) coincides well with the point in the template where a transition between a thick and well distinct layer 4 (A3b) and a thin and less distinct layer 4 (A3a) occurs in the Nissl volume (Fig. 4B). Moving further medially, the transition to area 4ab (which corresponds to the body and limb representations of the primary motor area; Burman et al., 2014a) coincides with the disappearance of any evidence of layer 4 in the sections through the Nissl volume.

To evaluate the spatial variability of areas across the 20 animals, we conducted an analysis based on the assignment of individual streamlines to areas (Fig. 5E). This analysis, based on 829 514 streamlines, shows that the majority of these could be attributed to a single (43%) or one of 2 adjacent (40%) areas, across the 20 individuals. However, some of the streamlines could belong to as many as four or five different areas (2.4% and 0.2%, respectively), thus revealing significant variability (a small group of 48 streamlines, <0.01% of the total, was assigned to 6 different areas). Interestingly, many of the high-variability streamlines concentrated in sectors which have expanded differentially in primate brain evolution, as previously shown, including the temporoparietal junction, ventrolateral prefrontal and cingulate regions (Chaplin et al., 2013).

The provided probability maps allow for estimation of the amount of variability in the spatial extent of a given cytoarchitectural area. In Figure 5F-H we demonstrate the distributions of streamlines associated with four example areas. This illustrates the fact that different areas vary in the extent to which they can be located based on stereotaxic coordinates alone. Some areas, such as A4ab and the middle temporal visual area (MT; Fig. 5F) are characterized by a core of high-confidence streamlines (0.95 isoprobability contour), surrounded by layers of streamlines with decreasing probabilities. In contrast, the stereotaxic coordinates of the lateral intraparietal (LIP) and primary somatosensory areas (A3b), are inherently uncertain (Fig. 5F-H), even though these areas can be defined with the same level of precision using myeloarchitectural criteria (Krubitzer and Kaas, 1990; Burman et al., 2008; Rosa et al., 2009).

### 3.4 Cortical thickness

The information derived from the streamline analysis can be used to provide accurate estimates of the cortical thickness at a specific point (Fig. 4D). When combined with the gray level profiles shown in Fig. 4A, this information can be valuable in designing experiments, for example in situations where electrode recordings or injection sites need to be directed to specific cytoarchitectural layers of the cortex. For this purpose, the template also includes data layers corresponding to the distance from the pial surface (Figs. S2A, S3E, and S4E).

The estimates of cortical thickness varied across the cortex (Fig. 6A, B). The pattern retains many features previously observed in analyses based on an individual marmoset brain (Atapour et al., 2019). For example, as expected (Hilgetag and Barbas, 2006), regions of highly convex cortex (e.g. those along the dorsal midline, or the ventrolateral prefrontal convexity) tend to be noticeably thicker than cortex on the adjacent “flat” surfaces, or concave regions lying along the fundus of a sulcus. This pattern can be noticed even within single areas, such as V1 (Fig. 6A). Superimposed on this variation related to the local geometry of the cortex are region-specific features (Fig. 6B), which reflects previous findings in the macaque monkey (Koo et al., 2012). For example, areas in the ventral posterior parietal cortex, rostral temporal cortex, prefrontal cortex and precuneus stand out as being relatively thick in comparison with nearby areas, despite being located in unconvoluted surfaces. Conversely, limbic areas (including those dorsal to the corpus callosum, as well as parahippocampal, perirhinal and orbitofrontal areas) appear as relatively thin. Furthermore, a clear trend towards increased thickness following the rostrocaudal axis (Cahalane et al., 2012; Atapour et al., 2019) could be observed (Fig. 6D, E).

**Figure 6:**
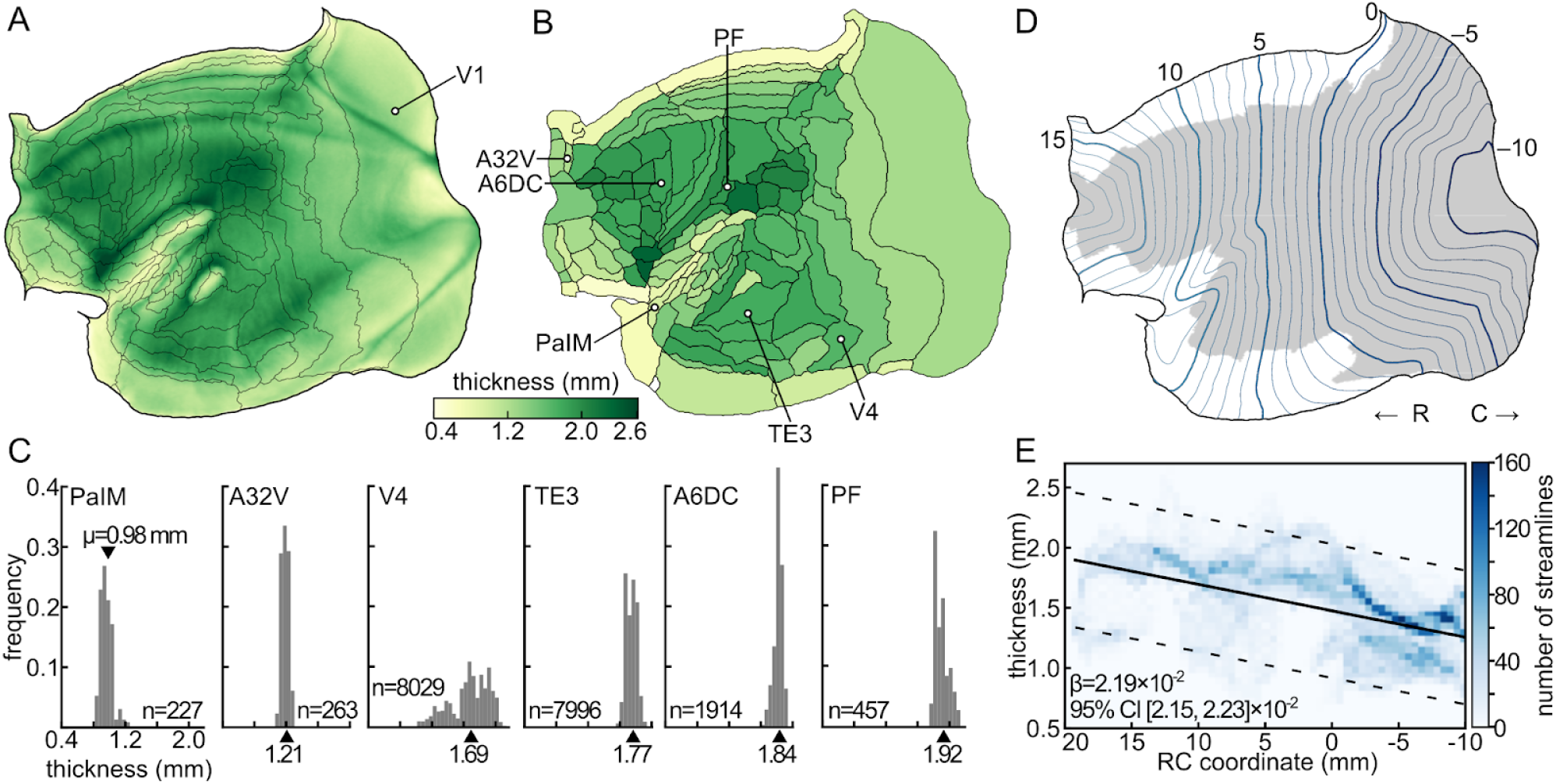
Cortical thickness. **A**: Map of the cortical thickness projected onto a flattened representation of the cortex. **B**: Mean cortical thickness of 116 cytoarchitectural areas of the marmoset brain. **C**: Distribution of the cortical thickness within six selected areas indicated in panel B. The *n* value denotes the number of streamlines selected for a particular cortical area while the black triangles indicate the average thickness as presented on panel B. **D**: Rostrocaudal stereotactic coordinates of the mid-thickness surface of the cortex projected onto the flatmap. The thick lines are plotted every 5 mm while the thin lines every millimeter. The gray shading indicates the combined extent of the isocortical areas that were used to characterize rostrocaudal gradient of cortical thickness (panel E); these were selected to maintain compatibility with previous studies (Cahalane et al., 2012; Atapour et al., 2019). **E**: Two-dimensional histogram of the cortical thickness and the rostrocaudal coordinate illustrating the gradient of cortical thickness along the rostrocaudal direction. The color of each bin represents the number of streamlines in a window of 0.05 mm (thickness) × 0.5 mm (rostrocaudal distance). The thick black line corresponds to the linear regression (F_1.30381_=1.15×10^4^, p<10^−4^, β=2.19×10^−2^, 95% CI [2.151, 2.231]×10^−2^) of the cortical thickness against the rostrocaudal coordinate while the dashed lines indicate the 95% prediction interval.

### 3.5 Laminar profiles

The streamline analysis also allowed the calculation of intensity profiles along columnar trajectories (Fig. 7A, C), which provides a way to quantify the macroscopic-level laminar differences observed in the template cortex (Figs 4, S3, S4). Although this analysis resulted in continuous estimates along the cortical surface, combined with the maximum likelihood estimates of area boundaries (Fig. 4B) it also allowed the calculation of Nissl image intensity profiles for all cytoarchitectural areas included in the template (see Fig. 7C for profiles of four example areas and the supplementary material for estimates for the remainder of the 116 areas). Variations in the Nissl image intensity are likely to reflect combined diversity in neuronal density and size, as well as the amount of intrinsic curvature of the cortex (Fig. 7D). Indeed, we found a strong relationship between the average Nissl image intensity of areas and their neuronal density (least-squares linear regression, F_1.116_= 42.80, p < 10^−4^, β = -2.02, 95% CI [-2.62, -1.40]), obtained using stereological analysis (Atapour et al., 2019) (Fig. 7E, F).

**Figure 7:**
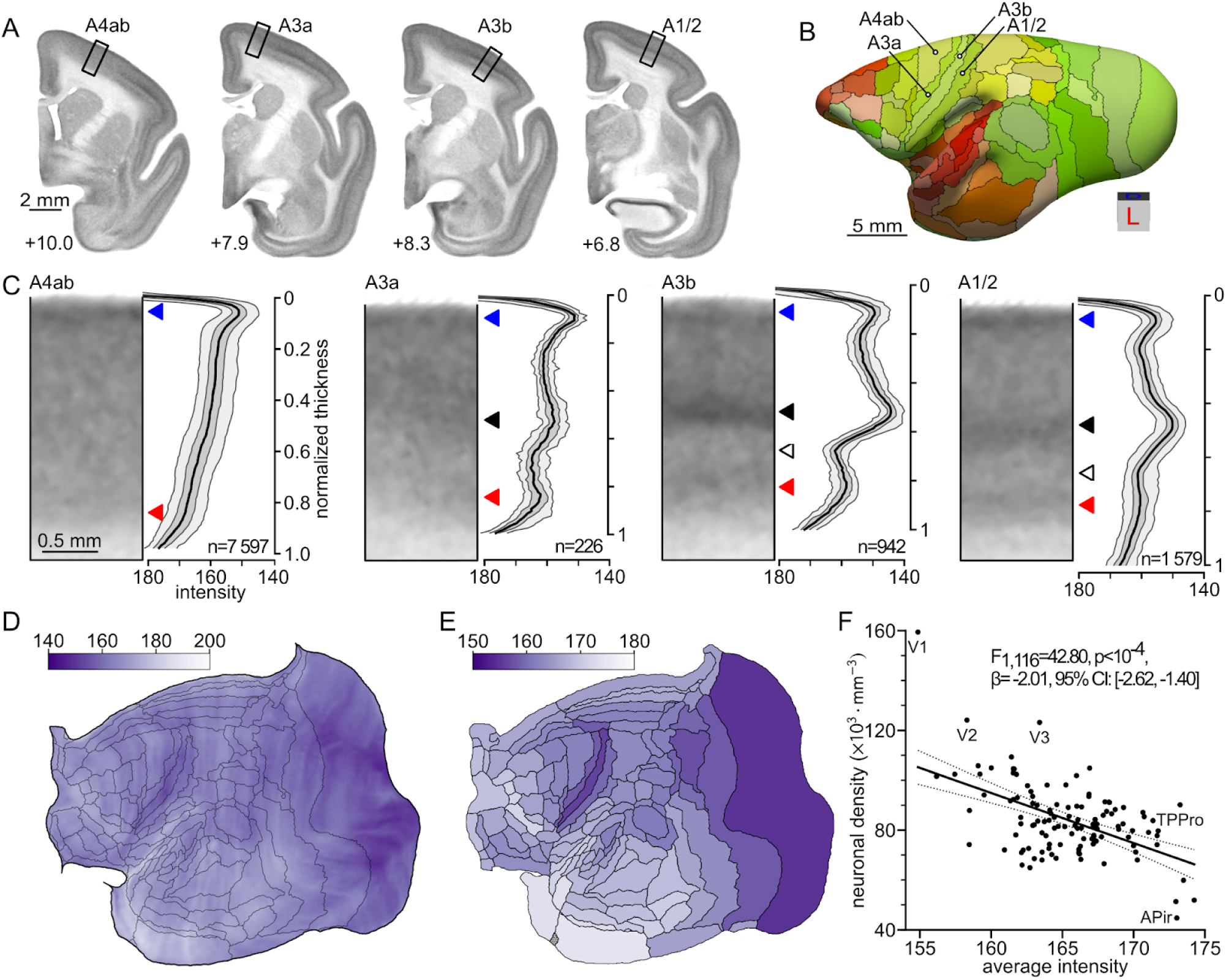
Laminar patterns of cortical areas. **A**: Coronal cross-sections through the template at four rostrocaudal levels (A +6.8 to A +10.0; numbers next to the sections) showing the location of samples belonging to four adjacent cortical areas (black rectangles). **B**: Location of the samples indicated on panel A, presented against the mid-thickness surface of the cortex. **C**: Magnified views of the cross-sections (grayscale images) of the four example areas presented in panel A next to the staining intensity profiles (vertical line plots), illustrating differences in the laminar patterns of the areas. The top of the grayscale image (normalized thickness equal to 0) corresponds to the pial surface, while the bottom (normalized thickness of 1) indicates the white matter boundary. The thick black line denotes the median intensity among all streamlines at a given level of the normalized thickness, the light gray bands represent 5th and 95th centiles of the intensities, and the bands in dark gray correspond to the lower and upper quartiles. Triangles of different colors denote various characteristic parts of the profiles: layer 2 (blue), layer 4 (black), layer 5 (while), and layer 6 (red). Values next to each cross-section correspond to the number of streamlines with confidence level ≥ 90%, which were used to generate the profiles. Results for all 116 areas are available as supplementary material (http://www.marmosetbrain.org/nencki_monash_template). **D**: Average intensity per streamline (Fig. 3B) visualized against the flatmap. The grayscale intensity values (values of the red channel of the RGB Nissl template) are presented in the shades of purple to resemble the tint of the Nissl-stained sections. **E**: Average intensity per cortical area. The staining intensity reflects an approximately rostrocaudal gradient of neuronal density, previously described in the primate cerebral cortex (Cahalane et al., 2012; Atapour et al., 2019). **F**: Inverse-proportional relation between the staining intensity and neuronal density, both averaged per cortical area. The relation indicates that the template image intensity provides an adequate proxy of the neuronal density. For a list of areas and their abbreviations see Table S2.

### 3.6 Interoperability with other brain atlases

Finally, we address the interoperability between the NM template and other current digital atlases of the marmoset cortex: the Paxinos et al. (2012) template converted into a three-dimensional image (Majka et al., 2016), the Brain/MINDS 3D marmoset brain atlas (Woodward et al., 2018) and the MBM 2.0 high-resolution MRI template (Liu et al., 2020). To enable the bidirectional traversal between different atlas spaces, three sets of affine and deformable spatial transformations were computed. This makes it possible, for instance, to warp the images of these three atlases into the stereotaxis space of the NM template (Fig. 8A-D), but also vice versa – to map the data layers of the present template into any of the three coregistered atlases.

**Figure 8:**
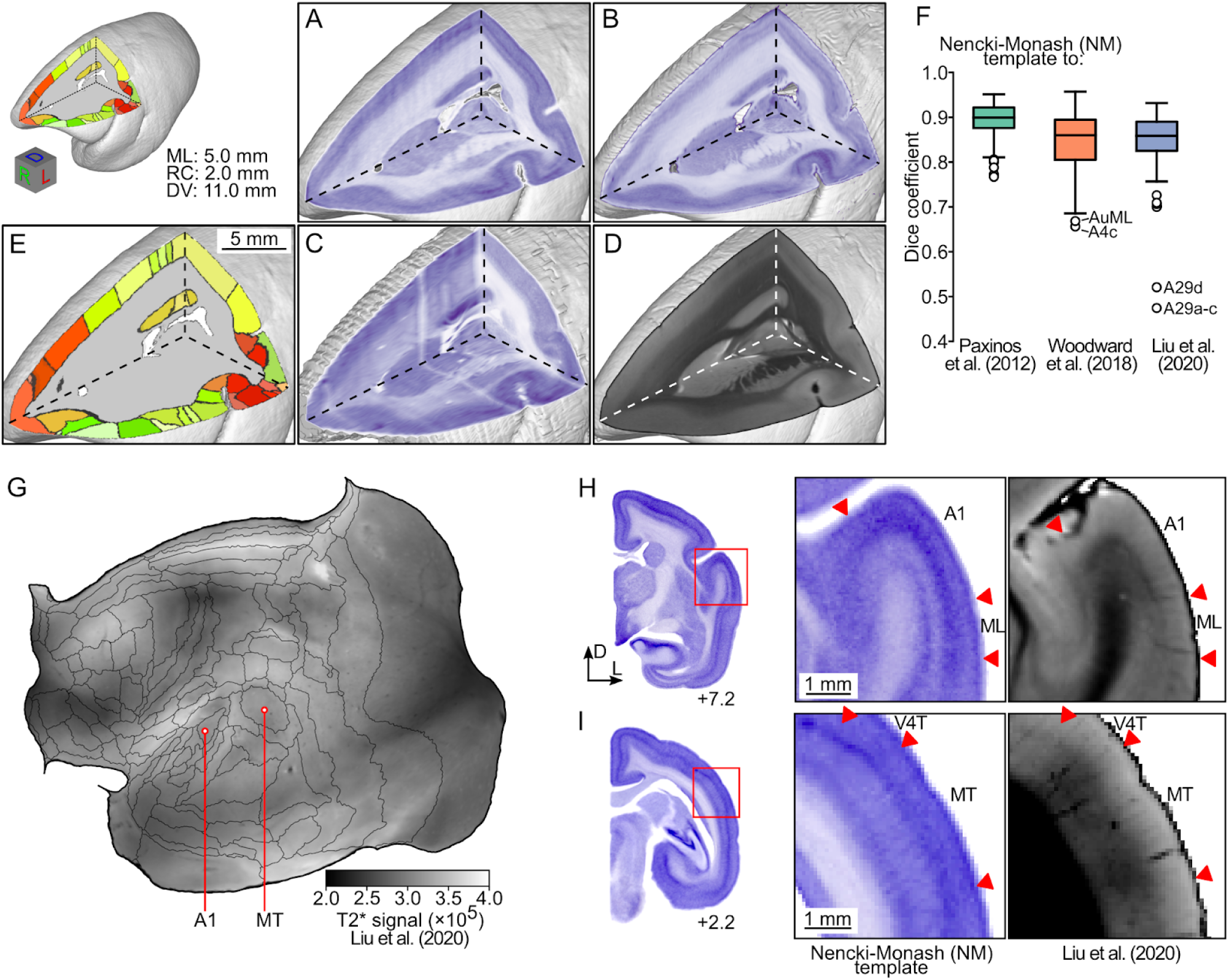
Atlas interoperability. **A**-**D:** Cross-sectional views of 3D reconstructions of the marmoset dorsolateral prefrontal cortex, comparing the histological appearance of this part of the brain in the NM template (**A**, present data), the Brain/MINDS marmoset brain atlas (**B**; Woodward et al., 2018), a volumetric reconstruction of the Paxinos et al., (2012) atlas (**C**; Majka et al., 2016), and the MBM atlas 2.0 (**D**; Liu et al., 2020). A-C are based on Nissl stain, and D on ex-vivo MRI (T2* contrast). The subdivision of the cortex into areas is illustrated in panel E, according to the present template. **F**: Registration accuracy expressed with the Dice coefficient. The box plots (centerline: median; box limits: upper and lower quartiles; whiskers: 1.5 × interquartile range; points: outliers) represent distributions of the values of the overlap between pairs of corresponding areas. **G**: T2* signal reported by Liu et al., (2020) registered to the NM template, presented using the flat map representation of the cortex. **H, I:** Comparisons of sections through the dorsal temporal lobe demonstrate the good correspondence between the borders of areas in the Nissl intensity image (middle panels, present data) and the coregistered T2* MRI (right panels, from the MBM 2.0 atlas). For a list of areas and their abbreviations see Table S2.

To determine the accuracy of the obtained spatial transformations, the cortical areas according to the NM segmentation were compared with those represented in the other atlases, after mapping them to the NM space. The assessment took advantage of the fact that the parcellation of all considered templates follows the Paxinos et al. (2012) criteria, thus enabling the direct calculations of the overlap between corresponding cortical areas, for instance, using the Dice coefficient statistic.

The benchmark showed that the coregistration procedure yielded accurate spatial transformations (Fig. 8F): The median Dice coefficient between the NM and the Paxinos et al. (2012) segmentations amounted to 0.90, with an interquartile range (IQR) of 0.04. For the two other templates, the accuracy was lower, but still high in absolute terms: median of 0.86 for each of the two other atlases, and IQRs of 0.09 and 0.07 (Brain/MINDS and MBM 2.0 templates, respectively). The spatial pattern of the coregistration accuracy was consistent for all three mappings (Fig. S6A-C). The computed alignment was highly precise for the majority of the 116 cortical areas, being particularly good for large areas, of regular and round shape (e.g. A8b, A4ab, PE, A8aV). On the other hand, smaller areas (e.g. A4c), those with an irregular shape (e.g. high elongation), or located in the parts of the cortex that are highly curved (e.g. A29a-c, A29d, IPro, and ProSt) tended to coregister less accurately. Three additional metrics were used to more comprehensively evaluate the spatial patterns of the coregistration accuracy (Fig. S6D-F): the Jaccard coefficient, together with the maximum and average Hausdorf distances (the latter two statistics evaluate the agreement between the perimeters of cortical areas). All metrics yielded results consistent with those obtained by measuring the overlap using the Dice coefficient. The evaluation of the coregistration accuracy enables the informed application of the mapping between the NM and the other templates (i.e. taking into account the fact that in some parts of the cortex the mapping is more accurate than in others).

Utilizing the obtained spatial transformations, we have warped ultra-high resolution MR images of the MBM 2.0 atlas (Liu et al., 2020) into the space of the NM template. With these additional layers, the NM template provides an accurate multimodal view of the entire brain incorporating both cytoarchitecture and MR imaging modalities (Figs. 8 and S7). For instance, the warped T2* image helps to examine the myelination level of the individual areas (Fig. 8G). The borders between the areas align well with the abrupt changes in the myelination level in specific areas known from the literature for high myelination, such as AuA1 or MT (Fig. 8H, I; Rosa and Elston 1998; de la Mothe et al., 2006). The transformation also allows merging diffusion MRI data into the NM template (Fig. S7), which not only provides extra image contrasts, such as fractional anisotropy and mean diffusivity, but also demonstrates fiber orientations of the white matter running underneath the cortex (Fig. S7C).

## 4. Discussion

We introduce an open access template of the marmoset cerebral cortex, which reflects the morphological average of 20 cerebral hemispheres reconstructed from histological sections using the procedure described by Majka et al. (2016, 2020). The primary modality of the template is a Nissl-stained volume, which can be virtually cross-sectioned in any plane. The main noticeable difference from the previously available histology-based templates (Hashikawa et al., 2015; Majka et al., 2016; Woodward et al., 2018) is its isotropic nature. The averaging resulted in suppression of the artifacts and noise inherent to the histological processing of the individual hemispheres used to generate the template, in particular those related to the separation between sections (Fig. S8A). The obtained mean morphology of the marmoset cortex shows differences in comparison with those of the individual brains which can be illustrated by comparing, for instance, coronal planes at corresponding AP levels in the Paxinos et al. (2012) atlas and the present template. In contrast with the Paxinos atlas, the Nencki-Monash template shows a prominent superior temporal sulcus, which both indicates that this morphological landmark is common among the adult marmoset population, and that its location in the stereotaxic space is stereotypical (Fig. S8B). Conversely, the intraparietal sulcus which can be prominent in individual brains (e.g. Paxinos et al., 2012), is barely discernible in the Nencki-Monash template (Fig. S8C).

### 4.1 Methodological considerations

The present atlas is based on a process that takes into account both the morphological features of the 20 individual brains, and the locations of a subset of cytoarchitectural areas. The decision to inform the registration process on histological boundaries recognizes the challenges involved in trying to summarize both the average cortical anatomy and its variability across a genetically diverse population, and the fact that borders of cortical areas do not necessarily follow gyrification patterns. This contrasts with the approach taken by most large-scale projects aimed at mapping cellular connectivity in mice, which focus on individuals of a specific strain, sex, and age.

The decision to incorporate labels corresponding to the borders of areas was based on our experience in interpreting patterns of connections in the marmoset cortex, where we found that the locations of tracer injection sites determined solely on the intensity-based registration to a template often resulted in misattribution to an adjacent area (Majka et al., 2020); however, the incorporation of area-based label maps to guide the registration mitigated this problem. At the same time, we found it impractical to delineate all 116 cortical areas in each of the reconstructed hemispheres. A key consideration here is the fact that areas differ markedly in terms of the precision of identification of their boundaries, even when multiple histological stains are used (Rosa and Tweedale, 2005; Paxinos et al., 2012). For example, whereas incorporating the boundaries of primary sensory areas always provides precise guidance to the configuration of the cortical mosaic in different hemispheres, the attempts to delineate single subdivisions of the ventral parietal cortex (PG, PFG, PF) or inferior temporal cortex (TE1, TE2, TE3) carry a much higher margin of error in most individual marmosets. We took the approach to delineate only areas that could be unambiguously defined. Thus, the area labels used in the registration process include single areas that could be sharply delimited (through the combined use of the Nissl, Gallyas and cytochrome oxidase stains) in a particular individual, as well as larger aggregations of areas that could be well distinguished from the neighboring cortex (e.g. the auditory core and belt areas as a whole, the combined extent of areas 13L and 13M, or the combined extent of all subdivisions of area TE). The process of calculation of the borders of other areas carries the assumptions that each marmoset brain has the same complement of cortical areas, and that the general spatial configuration of the areas relative to each other is more or less preserved across individuals (and both hemispheres). Although these assumptions are reasonable at this point, one must recognize that much still needs to be learned about the true extent of individual variability in the primate cortex. The present template can provide a means to investigate this issue further, for example by providing a “null hypothesis” against which future work using neuroimaging across large numbers of individuals can detect deviations from an expected pattern of cortical activation.

Are the brain hemispheres used to generate the template sufficient to accurately reflect the variability of the marmoset cerebral cortex? Other population based templates relied on a comparable (Hikishima et al., 2011) or smaller (Risser et al., 2019) numbers of individuals of similar sex and age, but did not incorporate histological validation of the area boundaries. Further, the morphological averaging was performed with a higher number of iterations (20) than recommended (Janke et al., 2015), which allowed to better emphasize cytoarchitectonic features and transitions between individual brain structures. Thus, the present atlas likely provides a good representation of the average morphology and variability of the cortex among young adult, captive-born marmosets. However, the template does not capture the changes associated with early development or senescence, which are likely to be significant (Missler et al., 1993; Uematsu et al., 2017; Sawiak et al., 2018), or the wider variability that could exist in wild populations, in which the extent of epigenetic changes associated with different postnatal experience is likely to be higher.

### 4.2 Atlas interoperability and future developments

The availability of the present template as an open resource provides additional layers of data for MRI-based templates, which enables cytoarchitectural correlations, thus establishing a link across imaging scales and modalities. The probabilistic mapping of cortical areas makes this resource particularly valuable for the purposes of planning experiments or analysing data. For example, registration of a structural MR image obtained from an individual to the present template can be used to direct tracer injections or electrode arrays to the coordinates most likely to belong to a specific area, or to assign levels of confidence to voxels revealed by functional imaging data. The feasibility of this approach is demonstrated by calculations of the transformations needed to register the present template to previously released high-resolution *ex vivo* MR images of the marmoset brain (Liu et al., 2020). This allowed us to correlate different structural features of various regions of the cortex, such as the T2* signal intensity and the Nissl image intensity, which, as the present study shows, is strongly correlated with neuronal density (Fig. 7F). Future registration to *in vivo* MRI-based templates (e.g. the IMPEC template; Risser et al., 2019) will also allow correction for possible distortions related to the histological processing, further increasing the precision of surgical approaches.

Whereas this interoperability opens the way for a multimodal parcellation of the marmoset brain, similar to that conducted for the human brain using structural and functional MRI data (Glasser et al., 2016), the present analyses also call attention to the challenges involved in this endeavour. For example, local variations related to cortical curvature were found within what is currently recognized as a single cytoarchitectural area (Fig. 6A). Combined with previous evidence of variations in cyto- and myeloarchitecture of single functional areas (e.g. the differences between parts of the primary motor cortex dedicated to control of facial, limb and axial musculature; Burman et al., 2008, 2014a), this indicates that attempts to obtain bias-free multimodal parcellations need to carefully consider both the geometry of the cortex and possible differences in the functional characteristics of the neurons within different topographic sectors of the same area (see also Palmer and Rosa, 2006). Functional connectivity data in the marmoset are becoming gradually available (e.g. Hori et al., 2020), adding a new dimension to such attempts. Data on neuronal connectivity that are not anchored to current descriptions of the boundaries of cytoarchitectural areas (Majka et al., 2020) and future work resulting in maps of the distribution of different neuronal subtypes (Krienen et al., 2019) can both enable neuroinformatics-based refinements of our understanding of the boundaries of cortical areas in this species.

Another desirable future extension of this work would be the probabilistic mapping of subcortical structures. The quality of the registration makes it possible to resolve such structures, thus providing an initial step in this direction. Here, again, the issue of precision in delineation is likely to be a factor: whereas some nuclei (e.g. the lateral geniculate nucleus, or the claustrum) can be delimited as sharply defined structures, others are characterized by gradual transitions. An approach based on larger aggregations, such as considering the mediodorsal and pulvinar thalamic complexes as wholes, is likely to facilitate the integration of subcortical data, based on future collaborative research.

## 5. Conclusions

We report on a new three-dimensional template of the marmoset cortex which represents the average of 20 young adult individuals of both sexes. This template is the outcome of a process that takes into account information not only about morphological features of the individual brains, but also about the borders of the most clearly defined cytoarchitectural areas. Together with the development of a streamline-based analysis procedure that incorporates biologically plausible assumptions, this method has resulted in a resource which allows direct estimates of the most likely coordinates of each cortical area, as well as quantification of the margins of error involved in assigning voxels to areas. The versatility of the present template is illustrated by registration to other marmoset brain atlases, including the MRI-based MBM 2.0 atlas, thus demonstrating the feasibility of cross-atlas interoperability. Moreover, the preservation of quantitative information about the laminar structure of the cortex can allow principled comparisons with analogous data from the human brain (Wagstyl et al., 2018).

In summary, the present template combines some of the main advantages of histology-based atlases (e.g. preservation of information about the cytoarchitectural structure, and direct estimates of the borders of areas) with features more commonly associated with MRI-based templates (e.g. isotropic nature of the dataset, and enablement of probabilistic analyses). The approach adopted here may be useful in the context of the future development of average brain templates that incorporate information about the variability of areas in other species for which it may be impractical to ensure homogeneity of the sample in terms of age, sex and genetic background. Given the rapid development of open access digital resources for the marmoset brain (Lin et al., 2019; Liu et al., 2020; Majka et al., 2020), and the availability of data on resting state functional connectivity (Hori et al., 2020), the present template may become an important contributor to studies that will lead to a new level of insight on the anatomical principles of organization of the primate brain.

## Data availability statement

Individual components of the Nencki–Monash template generated and analyzed in the current study are released under the terms of Creative Commons Attribution-ShareAlike 4.0 License and publicly available through the Marmoset Brain Connectivity Atlas portal (http://www.marmosetbrain.org/nencki_monash_template).

## Ethics statement

All input data used in the study are available via the Marmoset Brain Connectivity Atlas portal (http://www.marmosetbrain.org/) database. All experiments conducted for the purpose of creating this database conformed to the Australian Code of Practice for the Care and Use of Animals for Scientific Purposes, were approved by the Monash University Animal Experimentation Ethics Committee, and were described in detail in previous publications (Majka et al., 2016, 2020). Since the materials in this paper are digital images acquired from an openly-available online database, the work is exempt from institutional ethics review.

## Disclosure of competing interests or affirmative statement that there are none

Authors declare no conflict of interest.

## CRediT authorship contribution statement

*Conceptualization*: P.M., M.G.P.R.; *Methodology*: P.M., M.G.P.R.; *Software*: P.M., S.B., C.L, G.S.; *Validation*: P.M., M.G.P.R.; *Data Analysis*: P.M., S.B., J.M.C., N.J., C.L, G.S., K.H.W.; *Data Curation*: P.M., S.B., J.M.C., N.J., C.L, K.H.W., M.G.P.R.; *Writing*: P.M., M.G.P.R.; *Review & Editing*: All authors; Visualization: P.M., G.S., S.B.; *Supervision*: P.M., A.C.S., D.K.W., M.G.P.R.; *Project Administration*: P.M., D.K.W., M.G.P.R.; *Funding Acquisition*: P.M., A.C.S., D.K.W., M.G.P.R.

## Acknowledgements

The authors would like to acknowledge the contributions of Dr. Kathleen Burman, who performed several of the histological stains used in the generation of the template, and of Ms. Daria Malamanova and Dr. Ianina H. Wolkowicz, who performed the digitization of the histological slides. This work utilized multi-modal Australian ScienceS Imaging and Visualization Environment (MASSIVE) high performance computing infrastructure (https://www.massive.org.au/), and scanning of histological slides was performed by the Monash Histology Platform. Funding for the experiments and for the development of the resource was provided by International Neuroinformatics Coordinating Facility (INCF Seed Funding grant scheme), the National Centre for Research and Development (ERA-NET-NEURON/17/2017), the Australian Research Council (CE140100007), and the National Health and Medical Research Council (APP1122220).

