## Supplementary Materials for "Histology-based average template of the marmoset cortex with probabilistic localization of cytoarchitectural areas"

### Supplementary figures

**Figure S1**: Summary of the label maps delineated in experimental cases to improve the accuracy of registration-based segmentation of each reconstruction.

**Figure S2:** Continuation of Fig. 4.

**Figure S3 and S4:** Parasagittal (Fig. S3) and horizontal (Fig. S4) cross-sections at various mediolateral (Fig. S3) or dorsoventral (Fig. S4) levels illustrating the main layers of data constituting the template.

**Figure S5**: Improvement of the segmentation accuracy due to the incorporation of the label maps in the registration process.

**Figure S6**: Evaluation of the coregistration accuracy between the Nencki-Monash (NM) template and the three other atlases of the marmoset brain.

**Figure S7:** Further examples of interoperability between the NM template and an *ex vivo* MRI brain atlas.

**Figure S8**: Equivalent stereotaxic levels in different marmoset brain templates.

### Supplementary tables

**Table S1**: Summary of the label maps.

**Table S2**: Abbreviations of the names of cortical areas.

**Table S3**: Summary of the registration parameters.

### Template files

Individual components of the Nencki–Monash template generated and analyzed in the current study are released under the terms of Creative Commons Attribution-ShareAlike 4.0 License and publicly available through the Marmoset Brain Connectivity Atlas portal (http://www.marmosetbrain.org/nencki_monash_template).

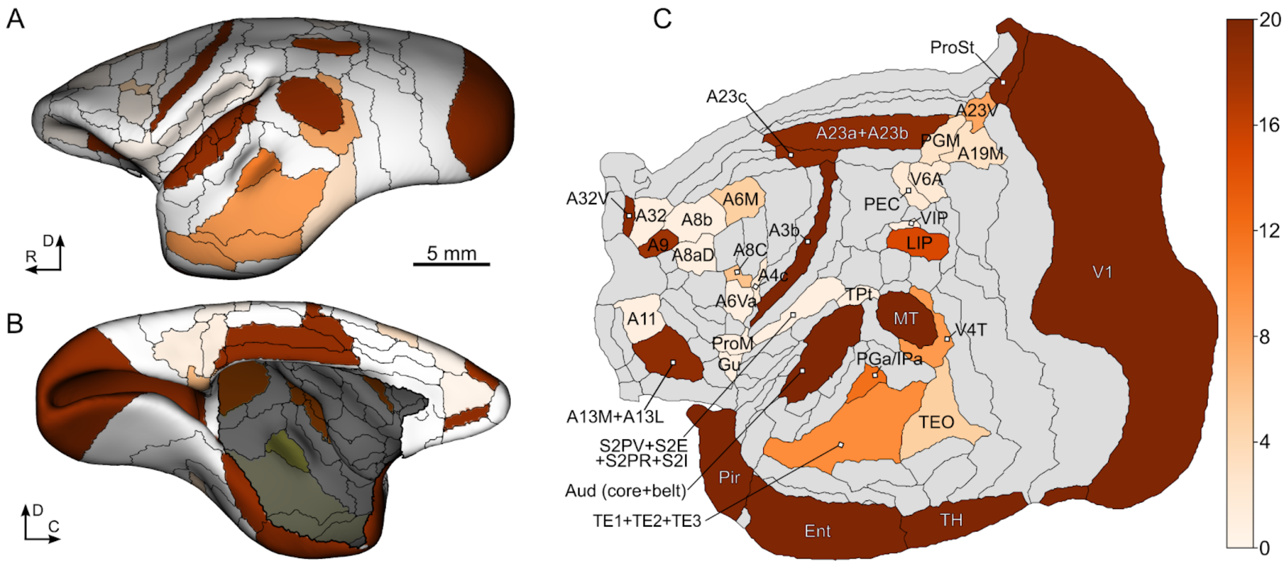


**Figure S1: Summary of the label maps delineated in experimental cases to improve the accuracy of registration-based segmentation of each reconstruction**. **A**, **B**: Medial and lateral views of the mid-thickness surface illustrating how many times, for the entire process of mapping all 20 reconstructions, individual cytoarchitectural areas were used to guide the registration process. **C**: Unfolded reconstruction of the cortex (left hemisphere representation; rostral to the left, medial to the top) presenting the same information. Note that eight label maps were drawn in each hemisphere: area 3b (A3b), auditory core and belt, Entorhinal cortex (Ent), Piriform cortex (Pir), prostriate area (ProS), temporal area TH, primary visual cortex (V1), middle temporal visual area (MT). Detailed information on areas outlined in each hemisphere is provided in Table S1.

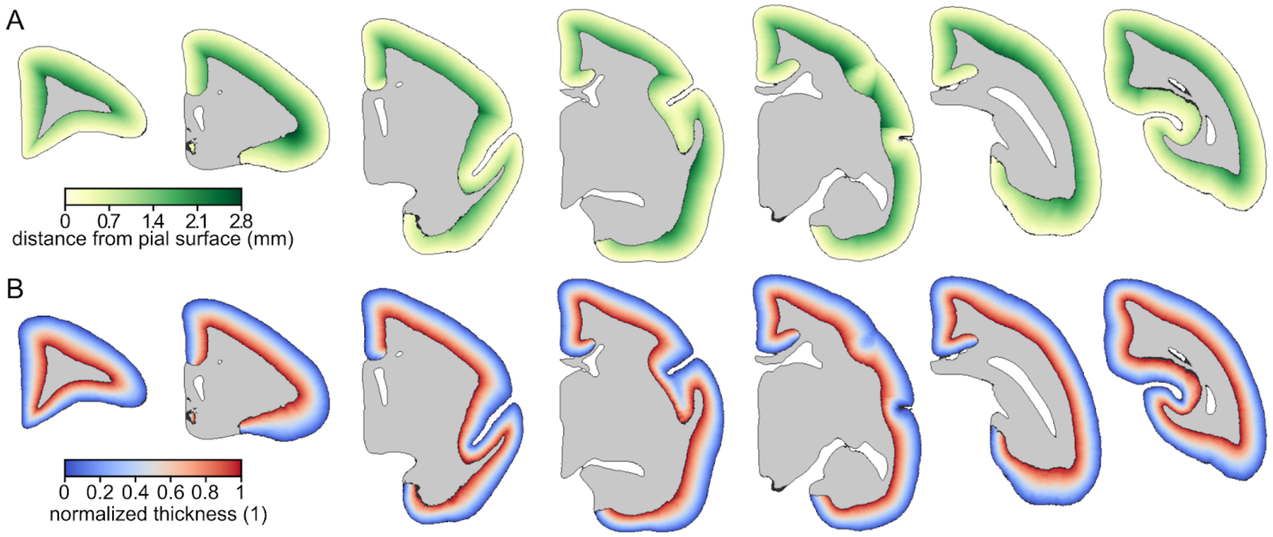


**Figure S2: Continuation of Fig. 4;** Coronal cross-sections at various anterior-posterior levels illustrating additional data layers provided with the template. **A**: Depth below the pial surface. **B**: Normalized thickness map (zero corresponds to the pial surface, one to the border between gray and white matter). All datasets are available for download from http://www.marmosetbrain.org/nencki_monash_template.

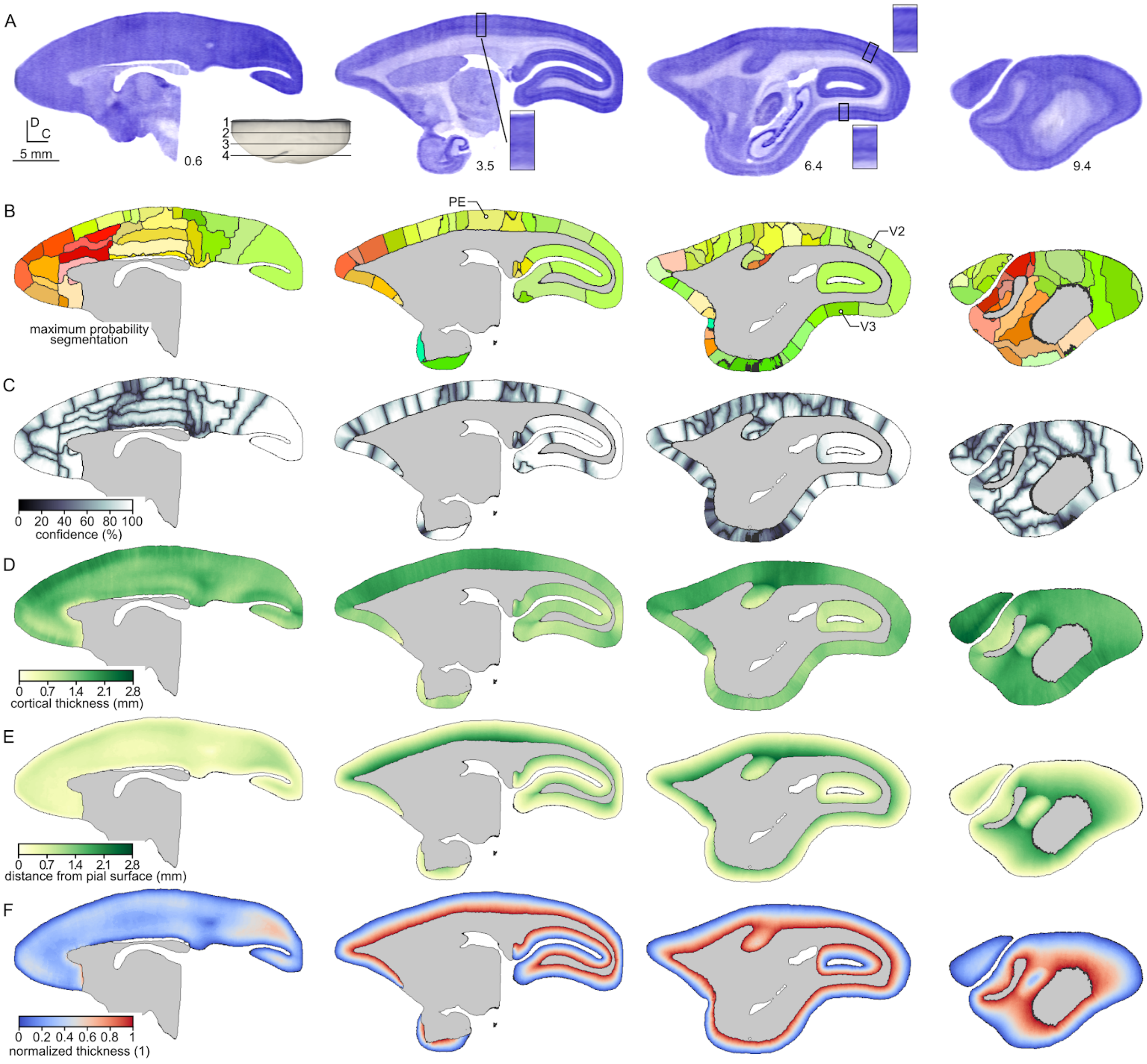


**Figure S3 and S4**: **Parasagittal (Fig. S3) and horizontal (Fig. S4) cross-sections at various mediolateral (Fig. S3) or dorsoventral (Fig. S4) levels illustrating the main layers of data constituting the template**. **A**: The template represented as a Nissl-stained volume. The magnified regions illustrate the laminar characteristics of selected cytoarchitectural areas labelled in panel B. Values next to each cross-section denote mediolateral (Fig. S3) or dorsoventral (Fig. S4) coordinates of the planes, which are also presented against the schematic views of the template (the gray, 3D model). **B**: Maximum-probability segmentation of the cerebral cortex into 116 areas created by selecting, for each cortical profile, the most probable label along a streamline (Fig. 3C, D). The black contours represent the borders of individual areas and are drawn for the clarity of the visualization. **C**: Segmentation confidence level map. **D**: Cortical thickness map. **E**: Depth below the pial surface. **F**: Normalized thickness map in which zero corresponds to the pial surface while one to the border between gray and white matter. All datasets are available for download from http://www.marmosetbrain.org/nencki_monash_template.

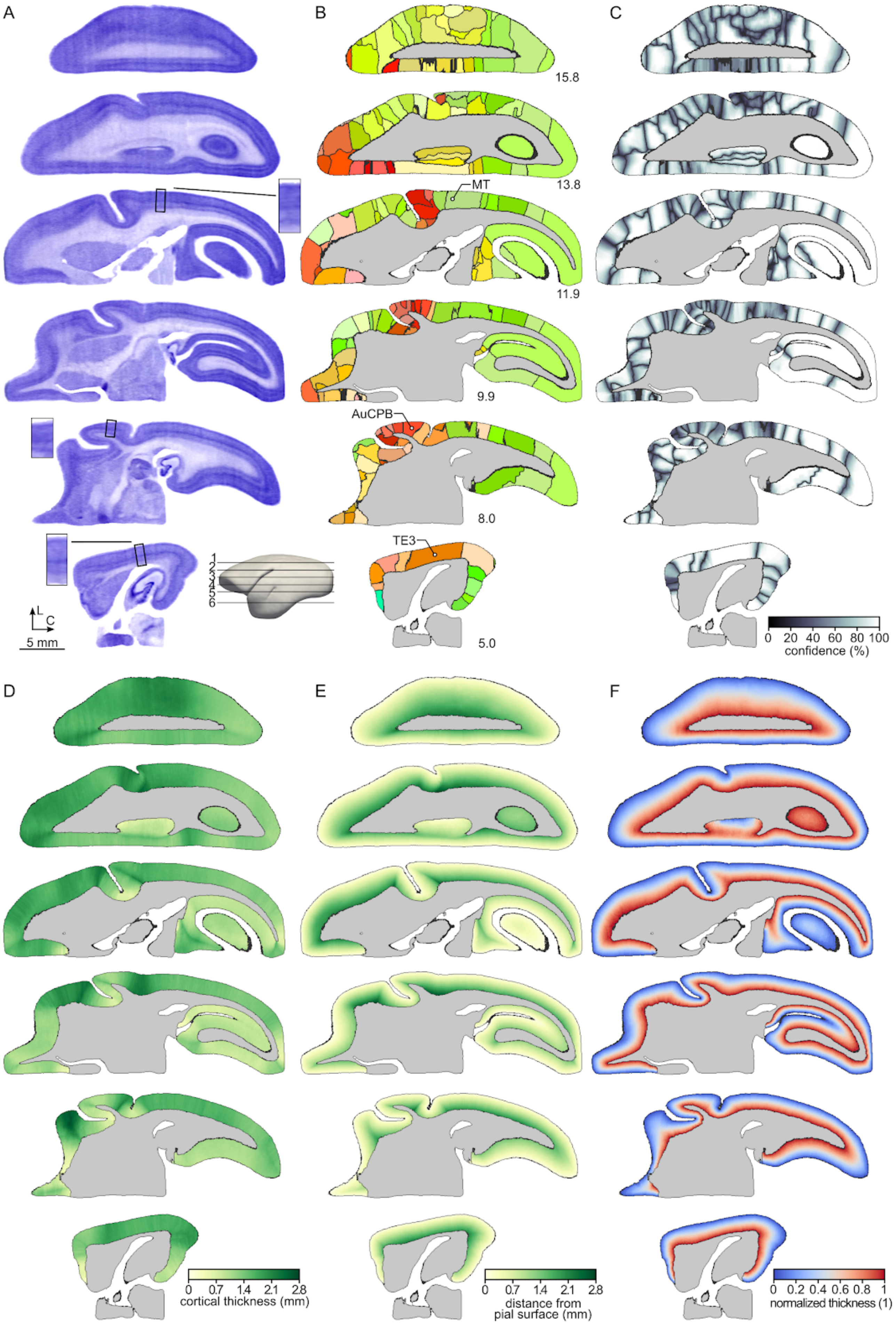
A
B
C
15.8
13.8
MT
11.9
9.9
AuCPB
8.0
TE3
5.0
1
2
3
4
5
6
L
C
5 mm
0 20 40 60 80 100
confidence (%)
D
E
F
0 0.7 1.4 2.1 2.8
cortical thickness (mm)
0 0.7 1.4 2.1 2.8
distance from
pial surface (mm)
0 0.2 0.4 0.6 0.8 1
normalized thickness (1)

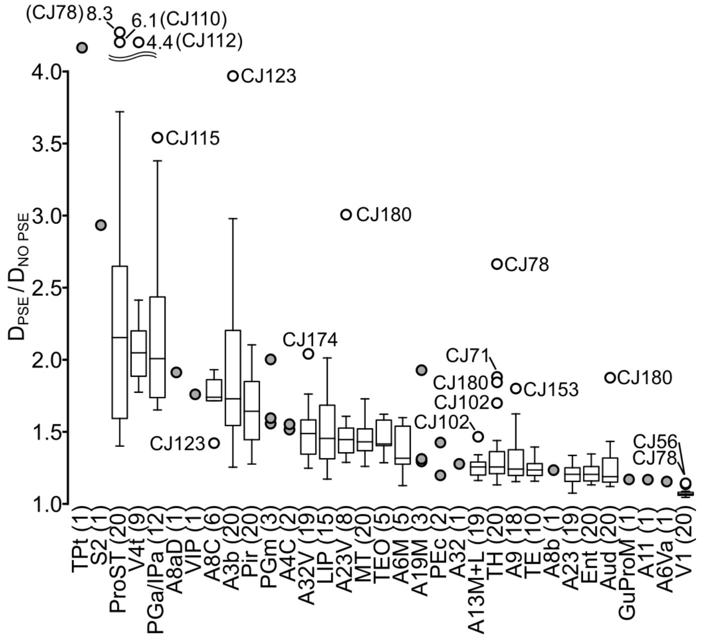

**Figure S5: Improvement of the segmentation accuracy due to the incorporation of the label maps in the registration process.** Comparison of the overlap between the outlines of selected cortical areas drawn manually on individual experimental cases and those of Paxinos et al. (2012) atlas after coregistration to the atlas *with* and *without* the use of label maps. $D_{PSE}$ denotes value of Dice coefficient obtained when the label maps were used to guide the registration, $D_{NO\ PSE}$ indicates value of Dice coefficient obtained when the mapping was performed without the use of the label maps. For all examined areas, the use of label maps increased the segmentation accuracy. The improvement was particularly noticeable for smaller areas (e.g. prostriata, PGa/IPa), with complex morphology (e.g. S2, A3b) or those located deep in sulci (e.g. TPt). Areas outlined in four or more cases are presented using box plots (center line: median; box limits: upper and lower quartiles; whiskers: 1.5 × interquartile range; annotated points: outliers), while those outlined in less than four cases are plotted with gray points. Source data is available in Table S1; for a list of areas and their abbreviations see Table S2.

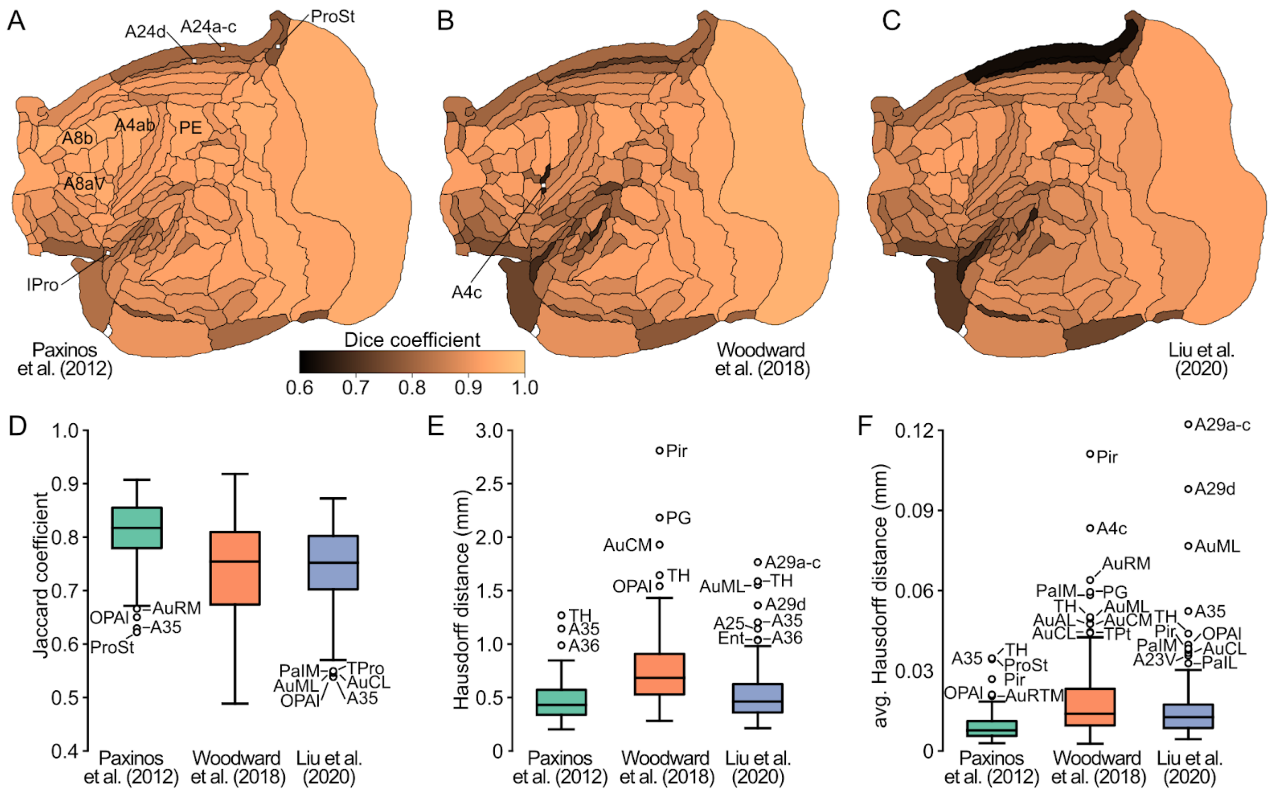


**Figure S6: Evaluation of the coregistration accuracy between the Nencki-Monash (NM) template and the three other atlases of the marmoset brain. A-C**: Values of the Dice coefficient measured between the segmentation of the NM template and segmentations of the other templates visualized on the flat map to illustrate the spatial pattern of the mapping accuracy. The pattern is consistent in all three cases: large areas of regular and round shape generally coregister better (e.g. A8b, A4ab, PE, A8aV) than those of irregular shape, smaller, elongated or located in the parts of the cortex that are highly curved (e.g. A29a-c, A29d, A4c, IPro). **D-F**: Evaluation of the coregistration accuracy using three additional statistics: Jaccard coefficient (D), Hausdorff distance (E), and average Hausdorff distance (F) for a more comprehensive overview of the obtained mappings. The results remain in agreement with the Dice coefficient (A-C, Fig. 8F). For a list of areas and their abbreviations see Table S2.

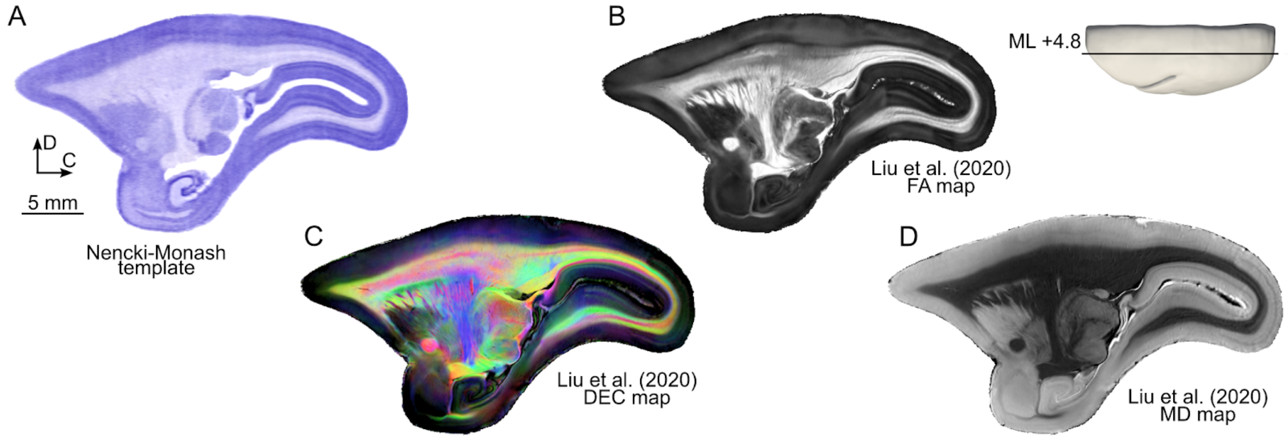


**Figure S7**: **Further examples of interoperability between the NM template and an *ex-vivo* MRI brain atlas.** Comparison of equivalent parasagittal cross-sections (ML +4.8 mm) of the Nencki-Monash template (**A**) with corresponding cross-sections of the diffusion MRI dataset of the MBM 2.0 atlas (Liu et. al., 2020; https://marmosetbrainmapping.org/) mapped to the NM template space **B**: Directionally encoded color (DEC) maps illustrating the fiber tracts passing in the rostrocaudal (shades of green), mediolateral (shades of red), dorsoventral (shades of blue), and diagonal (combination of the basic colors) directions. **C**: Fractional anisotropy (FA) map **D**: Mean diffusivity (MD) map. The datasets presented in this figure are available for download from http://www.marmosetbrain.org/nencki_monash_template.

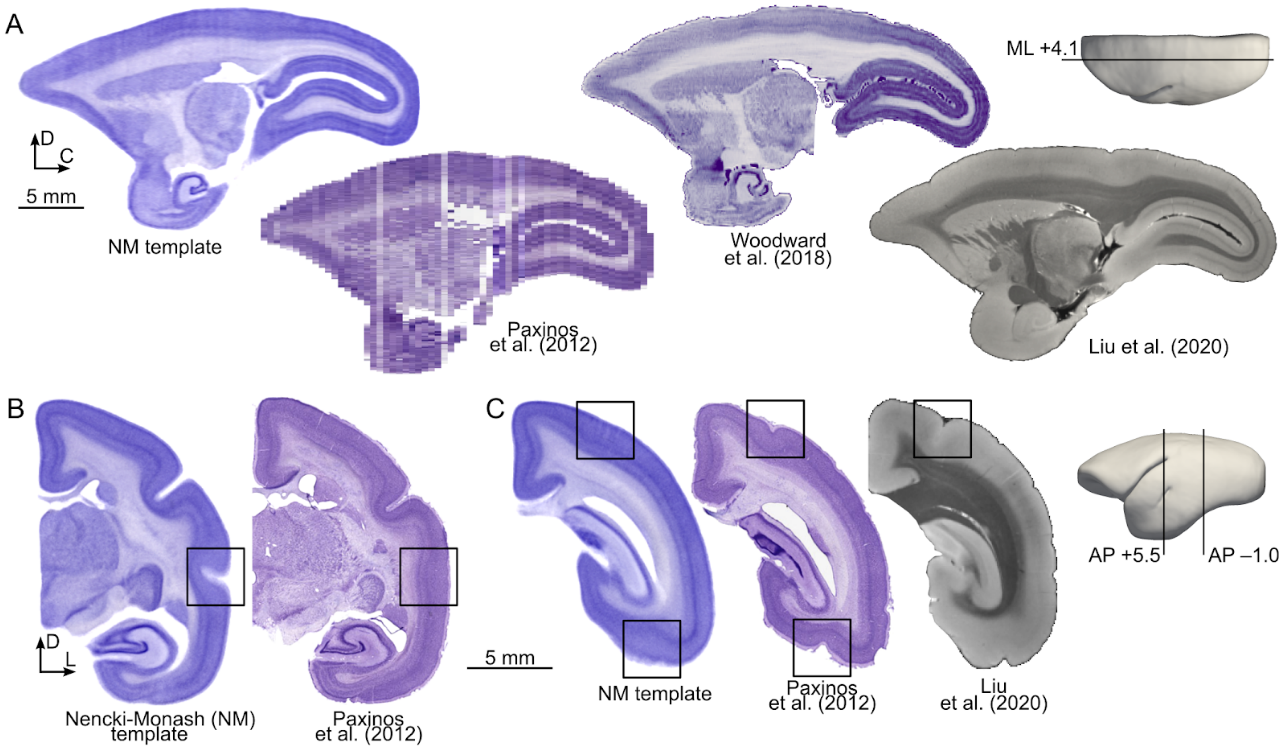


**Figure S8: Equivalent stereotaxic levels in different marmoset brain templates. A**: The averaging procedure employed in the present study mitigates histological artifacts that characterize templates generated from coronal (Paxinos et al., 2012) or horizontal (Woodward et al., 2018) sections. For comparison, a section obtained from an isotropic *ex vivo* MR image is illustrated (Liu et al., 2020). **B**: Comparison of equivalent coronal sections (AP +5.5 mm) demonstrate a robust superior temporal sulcus (box) in the morphological average, but not in the brain studied by Paxinos et al. (2012). **C**: At level AP -1.0 mm, the intraparietal sulcus is visible in individual brains (Paxinos et al., 2012, Liu et al., 2020), but not in the morphological average (NM template), indicating significant variability in the location of this landmark.

### Table S1: Summary of label maps

The numbers indicate the improvement of registration accuracy (measured with the Dice coefficient) when a label map was applied.
For instance, a value of 1.31 means that the dice coefficient was 1.31 times higher when a label map was used in comparison when it was not.

| Area Name | CJ181 | CJ123 | CJ122 | CJ116 | CJ153 | CJ164 | CJ174 | CJ108 | CJ157 | CJ114 | CJ115 | CJ78 | CJ180 | CJ102 | CJ112 | CJ111 | CJ81 | CJ56 | CJ110 | CJ71 | Number of times outlined |
|---|---|---|---|---|---|---|---|---|---|---|---|---|---|---|---|---|---|---|---|---|---|
| Area 11 of cortex | - | - | - | - | - | - | 1.17 | - | - | - | - | - | - | - | - | - | - | - | - | - | 1 |
| Area 13 of cortex lateral part | 1.27 | 1.29 | 1.20 | 1.32 | 1.17 | 1.18 | - | 1.31 | 1.21 | 1.34 | 1.23 | 1.29 | 1.20 | 1.47 | 1.28 | 1.16 | 1.26 | 1.26 | 1.19 | 1.23 | 19 |
| Area 13 of cortex medial part | 1.27 | 1.29 | 1.20 | 1.32 | 1.17 | 1.18 | - | 1.31 | 1.21 | 1.34 | 1.23 | 1.29 | 1.20 | 1.47 | 1.28 | 1.16 | 1.26 | 1.26 | 1.19 | 1.23 | 19 |
| Area 19 of cortex medial part | - | - | 1.31 | - | - | - | - | - | - | - | - | - | 1.93 | 1.29 | - | - | - | - | - | - | 3 |
| Area 23 of cortex ventral part | - | 1.29 | - | 1.43 | 1.46 | - | - | - | - | 1.61 | 1.32 | - | 3.01 | 1.50 | - | - | - | - | - | 1.36 | 8 |
| Area 23a of cortex | - | 1.17 | 1.18 | 1.21 | 1.33 | 1.22 | 1.15 | 1.12 | 1.26 | 1.07 | 1.13 | 1.21 | 1.24 | 1.27 | 1.20 | 1.14 | 1.16 | 1.34 | 1.21 | 1.27 | 19 |
| Area 23b of cortex | - | 1.17 | 1.18 | 1.21 | 1.33 | 1.22 | 1.15 | 1.12 | 1.26 | 1.07 | 1.13 | 1.21 | 1.24 | 1.27 | 1.20 | 1.14 | 1.16 | 1.34 | 1.21 | 1.27 | 19 |
| Area 23c of cortex | - | 1.17 | 1.18 | 1.21 | 1.33 | 1.22 | 1.15 | 1.12 | 1.26 | 1.07 | 1.13 | 1.21 | 1.24 | 1.27 | 1.20 | 1.14 | 1.16 | 1.34 | 1.21 | 1.27 | 19 |
| Area 32 of cortex | - | - | - | - | - | - | - | - | - | - | - | - | - | - | - | - | - | - | - | 1.28 | 1 |
| Area 32 of cortex ventral part | 1.29 | 1.59 | 1.41 | 1.49 | 1.25 | 1.37 | 2.04 | 1.58 | 1.55 | 1.43 | 1.70 | 1.76 | 1.31 | 1.31 | 1.52 | 1.52 | 1.32 | 1.59 | 1.39 | - | 19 |
| Area 3b of cortex (somatosensory) | 1.59 | 3.97 | 1.64 | 1.93 | 2.98 | 1.61 | 1.46 | 1.93 | 1.53 | 1.33 | 1.25 | 1.82 | 2.40 | 1.55 | 1.55 | 1.48 | 1.99 | 2.21 | 2.20 | 2.37 | 20 |
| Area 4 of cortex part c (primary motor) | - | - | - | - | - | - | - | - | - | - | 1.52 | - | - | - | - | 1.55 | - | - | - | - | 2 |
| Area 6 of cortex medial (supplementary motor) part | - | - | - | - | 1.13 | - | - | - | - | - | - | - | 1.60 | 1.54 | - | 1.28 | 1.32 | - | - | - | 5 |
| Area 6 of cortex ventral part a | 1.16 | - | - | - | - | - | - | - | - | - | - | - | - | - | - | - | - | - | - | - | 1 |
| Area 8 of cortex caudal part | - | 1.42 | - | 1.72 | 1.93 | - | 1.71 | - | - | - | - | - | - | - | - | 1.76 | - | 1.90 | - | - | 6 |
| Area 8a of cortex dorsal part | - | - | - | - | - | - | - | - | - | - | - | - | - | - | - | - | - | 1.91 | - | - | 1 |
| Area 8b of cortex | - | - | - | - | - | - | - | - | - | - | - | - | - | - | - | - | - | - | - | 1.23 | 1 |
| Area 9 of cortex | - | 1.19 | 1.24 | 1.15 | 1.80 | 1.62 | 1.27 | 1.19 | 1.24 | 1.23 | 1.43 | 1.41 | 1.21 | 1.17 | 1.22 | 1.25 | 1.26 | 1.49 | 1.18 | - | 18 |
| Auditory cortex primary area | 1.17 | 1.24 | 1.15 | 1.31 | 1.14 | 1.14 | 1.15 | 1.39 | 1.21 | 1.25 | 1.35 | 1.17 | 1.88 | 1.15 | 1.17 | 1.17 | 1.12 | 1.43 | 1.27 | 1.42 | 20 |
| Auditory cortex anterolateral area | 1.17 | 1.24 | 1.15 | 1.31 | 1.14 | 1.14 | 1.15 | 1.39 | 1.21 | 1.25 | 1.35 | 1.17 | 1.88 | 1.15 | 1.17 | 1.17 | 1.12 | 1.43 | 1.27 | 1.42 | 20 |
| Auditory cortex caudolateral area | 1.17 | 1.24 | 1.15 | 1.31 | 1.14 | 1.14 | 1.15 | 1.39 | 1.21 | 1.25 | 1.35 | 1.17 | 1.88 | 1.15 | 1.17 | 1.17 | 1.12 | 1.43 | 1.27 | 1.42 | 20 |
| Auditory cortex caudomedial area | 1.17 | 1.24 | 1.15 | 1.31 | 1.14 | 1.14 | 1.15 | 1.39 | 1.21 | 1.25 | 1.35 | 1.17 | 1.88 | 1.15 | 1.17 | 1.17 | 1.12 | 1.43 | 1.27 | 1.42 | 20 |
| Auditory cortex middle lateral area | 1.17 | 1.24 | 1.15 | 1.31 | 1.14 | 1.14 | 1.15 | 1.39 | 1.21 | 1.25 | 1.35 | 1.17 | 1.88 | 1.15 | 1.17 | 1.17 | 1.12 | 1.43 | 1.27 | 1.42 | 20 |
| Auditory cortex rostral area | 1.17 | 1.24 | 1.15 | 1.31 | 1.14 | 1.14 | 1.15 | 1.39 | 1.21 | 1.25 | 1.35 | 1.17 | 1.88 | 1.15 | 1.17 | 1.17 | 1.12 | 1.43 | 1.27 | 1.42 | 20 |
| Auditory cortex rostromedial area | 1.17 | 1.24 | 1.15 | 1.31 | 1.14 | 1.14 | 1.15 | 1.39 | 1.21 | 1.25 | 1.35 | 1.17 | 1.88 | 1.15 | 1.17 | 1.17 | 1.12 | 1.43 | 1.27 | 1.42 | 20 |
| Auditory cortex rostrotemporal | 1.17 | 1.24 | 1.15 | 1.31 | 1.14 | 1.14 | 1.15 | 1.39 | 1.21 | 1.25 | 1.35 | 1.17 | 1.88 | 1.15 | 1.17 | 1.17 | 1.12 | 1.43 | 1.27 | 1.42 | 20 |
| Auditory cortex rostrotemporal lateral area | 1.17 | 1.24 | 1.15 | 1.31 | 1.14 | 1.14 | 1.15 | 1.39 | 1.21 | 1.25 | 1.35 | 1.17 | 1.88 | 1.15 | 1.17 | 1.17 | 1.12 | 1.43 | 1.27 | 1.42 | 20 |
| Auditory cortex rostrotemporal medial area | 1.17 | 1.24 | 1.15 | 1.31 | 1.14 | 1.14 | 1.15 | 1.39 | 1.21 | 1.25 | 1.35 | 1.17 | 1.88 | 1.15 | 1.17 | 1.17 | 1.12 | 1.43 | 1.27 | 1.42 | 20 |
| Entorhinal cortex | 1.28 | 1.13 | 1.14 | 1.16 | 1.22 | 1.17 | 1.20 | 1.15 | 1.29 | 1.16 | 1.20 | 1.21 | 1.25 | 1.21 | 1.26 | 1.16 | 1.28 | 1.27 | 1.18 | 1.35 | 20 |
| Gustatory cortex | - | - | - | - | - | - | - | - | - | - | - | - | - | - | - | 1.17 | - | - | - | - | 1 |
| Lateral intraparietal area of cortex | - | 1.29 | - | 1.45 | 1.41 | 1.50 | 1.44 | 1.71 | 1.52 | 1.65 | 1.17 | - | 1.78 | 1.29 | 1.90 | - | - | 2.01 | 1.27 | 1.34 | 15 |
| Parietal area PE caudal part | - | - | - | - | - | - | - | - | - | 1.20 | - | - | - | - | - | - | 1.43 | - | - | - | 2 |
| Parietal area PG medial part (cortex) | - | 1.59 | - | - | 1.56 | - | - | - | - | - | - | 2.00 | - | - | - | - | - | - | - | - | 3 |
| Parietal areas PGa and IPa | - | 2.57 | 2.39 | - | 1.80 | 1.97 | 1.68 | 3.38 | 1.65 | 1.71 | 3.54 | - | - | - | 1.75 | - | - | 2.05 | 2.13 | - | 12 |
| Piriform cortex | 1.55 | 1.49 | 1.41 | 1.37 | 1.89 | 2.10 | 1.90 | 1.28 | 1.98 | 1.43 | 1.83 | 1.72 | 1.68 | 1.52 | 2.04 | 1.60 | 1.45 | 1.83 | 1.36 | 1.77 | 20 |
| Proisocortical motor region (precentral opercular cortex) | - | - | - | - | - | - | - | - | - | - | - | - | - | - | - | 1.17 | - | - | - | - | 1 |
| Prostriate area | 2.48 | 2.27 | 2.04 | 2.67 | 2.51 | 1.40 | 1.51 | 1.63 | 1.44 | 1.56 | 1.58 | 8.29 | 3.72 | 1.60 | 2.28 | 1.96 | 1.62 | 2.64 | 6.05 | 3.13 | 20 |
| Secondary somatosensory cortex external part | - | - | - | 2.93 | - | - | - | - | - | - | - | - | - | - | - | - | - | - | - | - | 1 |
| Secondary somatosensory cortex internal part | - | - | - | 2.93 | - | - | - | - | - | - | - | - | - | - | - | - | - | - | - | - | 1 |
| Secondary somatosensory cortex parietal rostral area | - | - | - | 2.93 | - | - | - | - | - | - | - | - | - | - | - | - | - | - | - | - | 1 |
| Secondary somatosensory cortex parietal ventral area | - | - | - | 2.93 | - | - | - | - | - | - | - | - | - | - | - | - | - | - | - | - | 1 |
| Temporal area TE1 (inferior temporal cortex) | - | 1.21 | - | 1.28 | - | - | - | 1.25 | 1.28 | 1.34 | 1.39 | - | 1.19 | 1.16 | - | - | - | 1.22 | - | 1.19 | 10 |
| Temporal area TE2 (inferior temporal cortex) | - | 1.21 | - | 1.28 | - | - | - | 1.25 | 1.28 | 1.34 | 1.39 | - | 1.19 | 1.16 | - | - | - | 1.22 | - | 1.19 | 10 |
| Temporal area TE3 (inferior temporal cortex) | - | 1.21 | - | 1.28 | - | - | - | 1.25 | 1.28 | 1.34 | 1.39 | - | 1.19 | 1.16 | - | - | - | 1.22 | - | 1.19 | 10 |
| Temporal area TE occipital part | - | - | - | - | - | - | 1.42 | 1.29 | 1.58 | - | - | - | - | - | 1.62 | - | - | - | 1.40 | - | 5 |
| Temporal area TH | 1.33 | 1.19 | 1.24 | 1.28 | 1.34 | 1.14 | 1.44 | 1.19 | 1.24 | 1.25 | 1.15 | 2.66 | 1.84 | 1.70 | 1.24 | 1.26 | 1.13 | 1.34 | 1.22 | 1.88 | 20 |
| Temporoparietal transitional area | - | - | - | - | - | - | 4.16 | - | - | - | - | - | - | - | - | - | - | - | - | - | 1 |
| Primary visual cortex | 1.08 | 1.08 | 1.07 | 1.07 | 1.09 | 1.04 | 1.06 | 1.05 | 1.06 | 1.06 | 1.06 | 1.14 | 1.06 | 1.07 | 1.08 | 1.06 | 1.06 | 1.14 | 1.06 | 1.08 | 20 |
| Visual area 4 transitional part (middle temporal crescent) | - | 1.89 | 1.78 | 1.87 | - | 2.20 | 2.41 | 2.05 | - | 2.12 | 1.89 | - | - | - | 4.42 | - | - | - | - | - | 9 |
| Visual area 5 (middle temporal area) | 1.46 | 1.43 | 1.28 | 1.42 | 1.36 | 1.34 | 1.58 | 1.53 | 1.26 | 1.42 | 1.52 | 1.34 | 1.72 | 1.47 | 1.73 | 1.46 | 1.41 | 1.60 | 1.37 | 1.43 | 20 |
| Visual area 6A (posterior parietal medial area) | - | - | - | - | - | - | - | - | - | 1.20 | - | - | - | - | - | - | 1.43 | - | - | - | 2 |
| Ventral intraparietal area of cortex | - | - | - | - | - | - | - | - | - | - | - | - | - | - | - | 1.76 | - | - | - | - | 1 |

Areas annotated with diagonal hatching were outlined as a single, merged region.

### Table S2: Abbreviations of the names of cortical areas

| | |
|---|---|
| A1-2 | areas 1 and 2 of cortex |
| A10 | area 10 of cortex |
| A11 | area 11 of cortex |
| A13L | area 13 of cortex lateral part |
| A13M | area 13 of cortex medial part |
| A13a | area 13a of cortex |
| A13b | area 13b of cortex |
| A14C | area 14 of cortex caudal part |
| A14R | area 14 of cortex rostral part |
| A19DI | area 19 of cortex dorsointermediate part |
| A19M | area 19 of cortex medial part |
| A23V | area 23 of cortex ventral part |
| A23a | area 23a of cortex |
| A23b | area 23b of cortex |
| A23c | area 23c of cortex |
| A24a | area 24a of cortex |
| A24b | area 24b of cortex |
| A24c | area 24c of cortex |
| A24d | area 24d of cortex |
| A25 | area 25 of cortex |
| A29a-c | area 29a-c of cortex |
| A29d | area 29d of cortex |
| A30 | area 30 of cortex |
| A31 | area 31 of cortex |
| A32 | area 32 of cortex |
| A32V | area 32 of cortex ventral part |
| A35 | area 35 of cortex |
| A36 | area 36 of cortex |
| A3a | area 3a of cortex (somatosensory) |
| A3b | area 3b of cortex (somatosensory) |
| A45 | area 45 of cortex |
| A46D | area 46 of cortex dorsal part |
| A46V | area 46 of cortex ventral part |
| A47L | area 47 (old 12) of cortex lateral part |
| A47M | area 47 (old 12) of cortex medial part |
| A47O | area 47 (old 12) of cortex orbital part |
| A4ab | area 4 of cortex parts a and b (primary motor) |
| A4c | area 4 of cortex part c (primary motor) |
| A6DC | area 6 of cortex dorsocaudal part |
| A6DR | area 6 of cortex dorsorostral part |
| A6M | area 6 of cortex medial (supplementary motor) part |
| A6Va | area 6 of cortex ventral part a |
| A6Vb | area 6 of cortex ventral part b |
| A8C | area 8 of cortex caudal part |
| A8aD | area 8a of cortex dorsal part |
| A8aV | area 8a of cortex ventral par |
| A8b | area 8b of cortex |
| A9 | area 9 of cortex |
| AI | agranular insular cortex |
| AIP | anterior intraparietal area of cortex |
| APir | amygdalopiriform transition area |
| AuA1 | auditory cortex primary area |
| AuAL | auditory cortex anterolateral area |
| AuCL | auditory cortex caudolateral area |
| AuCM | auditory cortex caudomedial area |
| AuCPB | auditory cortex caudal parabelt area |
| AuML | auditory cortex middle lateral area |
| AuR | auditory cortex rostral area |
| AuRM | auditory cortex rostromedial area |
| AuRPB | auditory cortex rostral parabelt |
| AuRT | auditory cortex rostrotemporal |
| AuRTL | auditory cortex rostrotemporal lateral area |
| AuRTM | auditory cortex rostrotemporal medial area |
| DI | dysgranular insular cortex |
| Ent | entorhinal cortex |
| FST | fundus of superior temporal sulcus area of cortex |
| GI | granular insular cortex |
| Gu | gustatory cortex |
| IPro | insular proisocortex |
| LIP | lateral intraparietal area of cortex |
| MIP | medial intraparietal area of cortex |
| MST | medial superior temporal area of cortex |
| OPAI | orbital periallocortex |
| OPro | orbital proisocortex |
| OPt | occipito-parietal transitional area of cortex |
| PE | parietal area PE |
| PEC | parietal area PE caudal part |
| PF | parietal area PF (cortex) |
| PFG | parietal area PFG (cortex) |
| PG | parietal area PG |
| PGM | parietal area PG medial part (cortex) |
| PGa/IPa | area PGa andIPa (fundus of superior temporal ventral area) |
| PaIL | parainsular cortex lateral part |
| PaIM | parainsular cortex medial part |
| Pir | piriform cortex |
| ProM | proisocortical motor region (precentral opercular cortex) |
| ProSt | prostriate area |
| ReI | retroinsular area (cortex) |
| S2E | secondary somatosensory cortex external part |
| S2I | secondary somatosensory cortex internal part |
| S2PR | secondary somatosensory cortex parietal rostral area |
| S2PV | secondary somatosensory cortex parietal ventral area |
| STR | superior temporal rostral area (cortex) |
| TE1 | temporal area TE1 (inferior temporal cortex) |
| TE2 | temporal area TE2 (inferior temporal cortex) |
| TE3 | temporal area TE3 (inferior temporal cortex) |
| TEO | temporal area TE occipital part |
| TF | temporal area TF |
| TFO | temporal area TF occipital part |
| TH | temporal area TH |
| TL | temporal area TL |
| TLO | temporal area TL occipital part |
| TPO | temporo-parieto-occipital association area (superior temporal polysensory cortex) |
| TPPro | temporopolar proisocortex |
| TPro | temporal proisocortex |
| TPt | temporoparietal transitional area |
| V1 | primary visual cortex |
| V2 | visual area 2 |
| V3 | visual area 3 (ventrolateral posterior area) |
| V3A | visual area 3A (dorsoanterior area) |
| V4 | visual area 4 (ventrolatereral anterior area) |
| V4T | visual area 4 transitional part (middle temporal crescent) |
| V5 / MT | visual area 5 (middle temporal area) |
| V6 | visual area 6 (dorsomedial area) |
| V6A | visual area 6A (posterior parietal medial area) |
| VIP | ventral intraparietal area of cortex |

**Table S3: Summary of the registration parameters**

| | | | Affine | | SyN | | | | CC | | PSE | |
|---|---|---|---|---|---|---|---|---|---|---|---|---|
| | Preprocessing | Registration resolution (µm) | Metric | Iterations | Gradient step | Regularization (displacement) | Regularization (velocity) | Iterations | Weight | Window size (vox) | Weight | Coverage |
| Individual cases (segmentation; Fig. 1) | median, 1 vox | 75 | MI | $10^3\times10^3\times10^3\times10^3\times10^3$ | 0.25 | 0.0 | 1.0 | $10^3\times10^3\times10^3\times10^3\times200$ | 1.0 | 4 | 1.0 | 100% |
| Morphological averaging (Fig. 2) | — | 100 | | N/A | | | | $200\times50\times50$ | | | N/A | |
| Registration to Paxinos et al. (2012) | | 100 | | | | | | | | | | |
| Registration to Woodward et al. (2018) | median, 1 vox | 75 | MI | $10^3\times10^3\times10^3\times10^3\times10^3$ | 0.25 | 0.0 | 1.0 | $10^3\times10^3\times10^3\times10^3\times200$ | 1.0 | 4 | 1.0 | 100% |
| Registration to Liu et al. (2020) | | 80 | | | | | | | | | | |

| | | |
|---|---|---|
| MI | Mutual Information | Avants et al., 2011 |
| CC | Cross Correlation | Avants et al., 2008 |
| PSE | Point set expectation | Avants et al., 2011 |

ANTs v. 2.1.0
